# Integration of Multiomic and Multi-phenotypic Data Identifies Biological Pathways Associated with Physical Fitness

**DOI:** 10.1101/2025.09.24.677381

**Authors:** Azar Alizadeh, John Graf, Matthew J. Misner, Andrew A. Burns, Fiona Ginty, Kevin J. O’Donovan, J. Kenneth Wickiser, Nicholas Barringer, Gregory Freisinger, Neil (Herm) Hermansen, J. Elizabeth McDonough, Brian Davis, Evelina R. Loghin, Christine Surrette, Peter Tu, Justin Welch, Oliver Boomhower, Ralf Lenigk, Rachel Sorrell, Tyler Hammond, Sara Peterson, Alison Caron, Leila Safazadeh, Chrystal Chadwick, Stephanie Stacey, James Jobin, Scott C Evans, Rui Xue, Gurvinder S. Khinda, Eric D. Williams, Swapnil Chhabra, Nhan Huynh, Ernest Fraenkel, Luca Marinelli

## Abstract

Unraveling the complex associations between human phenotypes and molecular pathways can pave the way to improved health and performance, but faces a fundamental challenge: the measurable genes, proteins, and metabolites vastly outnumber the participants in even the largest studies, yielding spurious correlations. To address this imbalance, we have developed a bioinformatic framework and computational approach (“PhenoMol”) to discover biological drivers of phenotypic characteristics that integrates all available phenotypic data predictive of outcomes and reduces multi-omic data dimensionality by generating “expression circuits” via graph theory constrained by prior biological knowledge of molecular interactions. We applied PhenoMol to analyze causal patterns and predict elite physical performance in a healthy cohort with deep physiological, physical, behavioral, cognitive, and molecular characterization. PhenoMol outperforms regression models based on equivalent analytic methodologies that do not employ network biology for dimensionality reduction. The PhenoMol software is provided for future studies.

## INTRODUCTION

Humans vary greatly in their ability to perform psychological and physical tasks, and a better understanding of the molecular basis for this variability would advance important medical, scientific and practical goals for improving life-long health and wellbeing. A number of studies have shown associations between epigenetic modifications and cognitive function and resiliency.^1, 2, 3^ The largest study to date with over 30,000 participants (from the CHARGE and COGENT consortia and UK Biobank) found 148 genome-wide significant independent loci associated with general cognitive function.^4^ Significant heterogeneity has also been reported in exercise response, where heritability has been shown to account for about 50% of cardiorespiratory fitness response variance^5^ and multiple on-going efforts are attempting to understand the molecular basis of response to exercise training.^6, 7, 8, 9, 10^ Beyond health and well-being applications, physical fitness, a key indicator of cardiovascular health^11^, is routinely used in both civilian and military selection processes to evaluate and enhance the potential and readiness of candidates for extreme roles. For example, professional sports teams hold events to assess physical readiness as part of selection pipelines (e.g., NFL, NBA and NHL Combines). Similarly, military corps have developed batteries of physical tests to rank and assess active-duty personnel (e.g., U.S. Army Combat Fitness Test (ACFT)^12^, US Navy Physical Readiness Test^13^). Furthermore, maintaining high levels of performance across many years is critical in military and athletic careers, and multiple studies are focused on uncovering the molecular basis of sustained fitness and resiliency.^14, 15, 16^

In addition to routine clinical laboratory biomarkers^17^, there is growing interest in using multi-omics analysis approaches to understand the molecular pathways that are modulated in response to physical stress and recovery, as well as to identify surrogate molecular markers of physical health and fitness to create a molecular portrait of an athlete.^18, 19^ For example, “sportomics” has focused on top level athletes from different sporting categories to identify metabolite changes during their recovery.^14,16^ The Athlome consortium, on the other hand, is comprised of several larger collaborative studies that are focused on DNA variants and genes associated with fitness and physical performance.^20^ Plasma proteomic profiling of healthy adults after 20 weeks of endurance training has identified hundreds of proteins associated with extracellular matrix remodeling, angiogenesis and iron homeostasis in the recovery period.^21^ Other studies have used metabolome^22^ or multi-parameter omics profiling to measure metabolic changes in response to activity.^23^ The shift towards omics-based assessment is driven by the fact that the effects of exercise are physiologically complex and multi-factorial, and no single measurement modality can realistically capture the activity of even one regulatory pathway. However, discovering statistically significant and predictive markers typically requires very large cohorts. For example, while Genome Wide Association Studies (GWAS) for discovery of genes associated with specific characteristics or risk factors^24^ often include tens of thousands of subjects, typical exercise intervention studies have much smaller numbers of participants. Multiple testing corrections combined with the typically modest effect sizes of these molecular measures severely limits the identification of significant predictors of high performance; furthermore, over-correction may be an issue when features are highly correlated.^25^

Computational algorithms and protocols for effectively combining physical/phenotype parameters with multi-parameter molecular markers (i.e., proteome, metabolome, genome, and clinical markers) to predict elite performance potential are still lacking.^18, 14^ To address this gap, we have developed a bioinformatic analysis pipeline (“PhenoMol”) that combines multiple layers of cellular and molecular biology data constrained by known molecular interactions in human biology in a graph theoretic framework. Network biology is a powerful constraint that enables the identification of candidate pathways (also referred to as expression circuits) using modest cohort sizes and amplifies effect sizes enabling discrimination of relevant phenotypes by individual molecules. It also provides robustness against the heterogeneity of human biology by establishing groups of molecules that act in concert to separate high-vs. low-performing subgroups, in contrast to individual markers.

We assembled a unique and highly curated dataset generated as part of the Measuring Biological Aptitude (MBA) program, which is sponsored by the Defense Advanced Research Program Agency and aims to identify, measure, and track personalized biomarkers related to training and performance for specialized military roles.^26^ Within the framework of the MBA program, we measured a broad battery of phenotypic and quotidian digital health measurements and performed longitudinal multi-omics profiling of blood components in a highly active cohort of cadets (*N*=86) from the United States Military Academy (USMA) at West Point. Participants were required to complete standardized physical fitness assessments, including the ACFT which is a composite measure of aerobic and anaerobic capacity, balance, flexibility, strength, agility, and perseverance.^12^ These cadets underwent up to five separate study segments in which they conducted intense cycling exercise (∼70% of VO_2_ max for 20 minutes). During each study segment, four separate intravenous blood samples were collected: one at baseline, and three after exercise (approximately <5, 10, and 30 minutes). Importantly, participants were randomly assigned to three periods of time throughout the day for their bike/blood draw events to account for variations in circadian rhythm. Samples were analyzed for untargeted metabolomics and proteomics, RNA sequencing, and DNA methylation. Hematological and immunophenotypic analyses, and targeted metabolic and cytokine panels, were also performed.

Using PhenoMol, we demonstrate that a network-based approach identifies functionally related molecules (expression circuits) that are predictive of participants’ physical fitness and performance. The network-based model used in this study is mechanistic, flexible (can include any -omics streams) and is designed for feature discovery in data sets with a sample of cohort size (N) much smaller than the numbers of molecular features (P).

## RESULTS

### Study Cohort and Design

To understand the biological pathways that drive exceptional physical performance, a broad range of phenotypic and molecular data was collected from 86 highly active subjects during a 3-month period (February-May 2021). The cohort was composed of 71 healthy male (age 21±1 years) and 15 healthy female (age 20±1 years) cadets from the USMA in West Point, NY, who were engaged in rigorous fitness and military training (∼1-5 hours/day). These cadets were required to complete standardized physical fitness assessments, including the ACFT, which consists of six separate athletic events: maximum deadlift (MDL), standing power throw (SPT), hand release pushups (HRP), sprint drag carry (SDC), leg tuck or plank (LTK), and two-mile run (2MR) (**Figure 1A**). The ACFT is scored on a scale of up to 600 points, with each of the events contributing a maximum of 100 points. The minimum score to pass the ACFT assessment is 360. The ACFT assessment in this study corresponds to the 2019 scoring system^27^, which mandated leg tuck and was scored on the same basis regardless of the age or sex of the participant.. The ACFT assessment was carried out towards the end of the three-month study, and the ACFT scores of 65 of the male cadets and 14 of the female cadets were available for this study.

**Figure 1.**
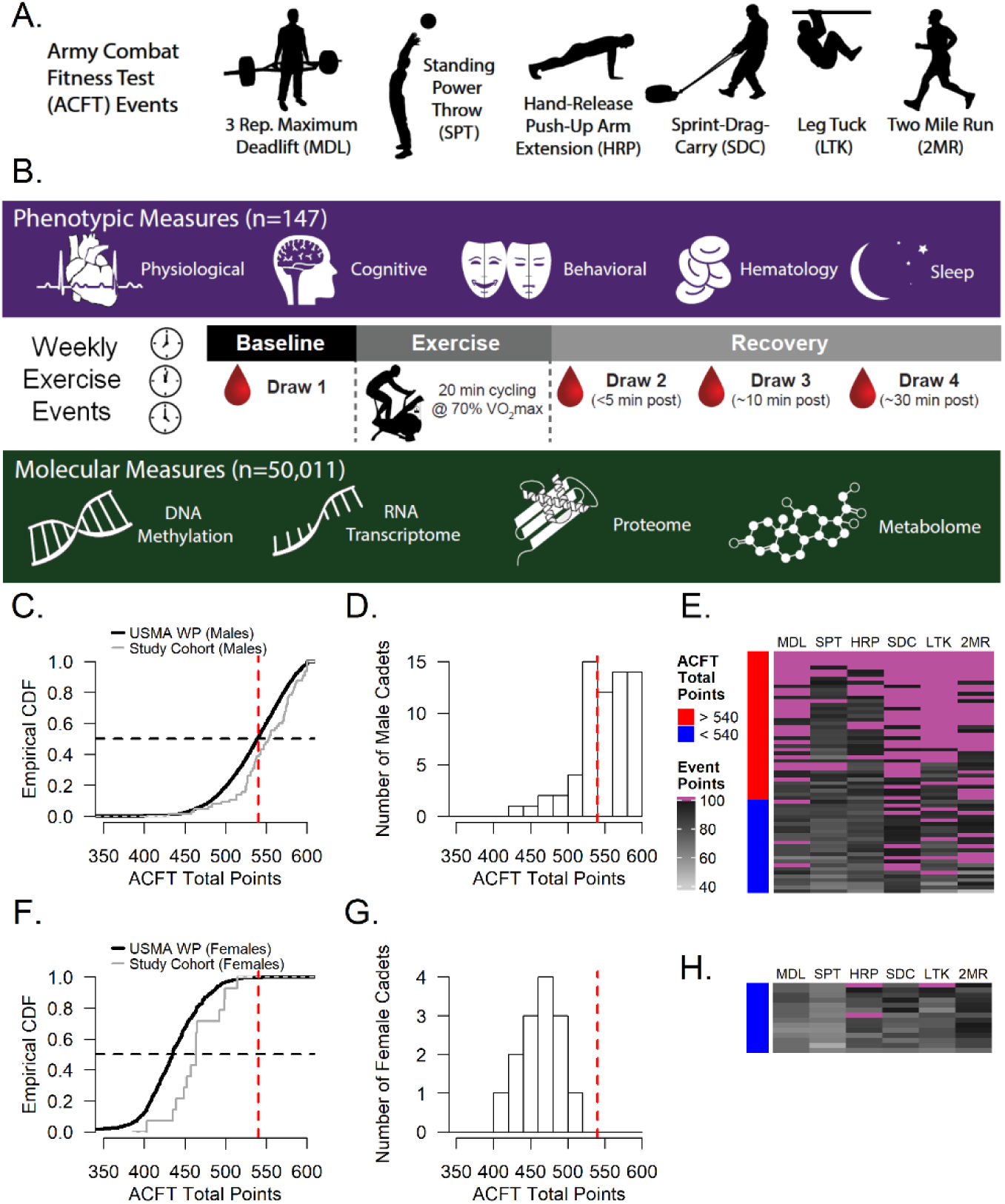
Measuring biological aptitude to achieve a 540+ Army Combat Fitness Test score. **A.** ACFT consists of six individual events (maximum deadlift (MDL), standing power throw (SPT), hand release pushups (HRP), sprint drag carry (SDC), leg tuck or plank (LTK), and two-mile run (2MR)) that are scored from 0 to 100 and summed into a total score, up to a maximum 600 points. **B.** Schematic representation of phenotypic measures including physiological, cognitive, behavior, hematology, and sleep. The exercise event was repeated weekly over five weeks along with random assignments to perform the exercise at three controlled time periods (morning, mid-day, late afternoon). Each exercise event consisted of a 20-minute strenuous cycling exercise at 70% VO2max with four blood draws, one at baseline, just prior to the bike exercise, and three draws approximately <5, 10, and 30 minutes after completion of the bike exercise. The types of molecular measurements (N=50,011) assayed from the drawn blood included DNA methylation of PBMCs, whole blood transcriptomics untargeted metabolomics and proteomics, and targeted biomarkers, including a cytokine panel. **C**. Cumulative distribution function (CDF) of the ACFT total points for the Spring 2021 USMA West Point male total population (thick black line, N=2,414); the study cohort of 65 male Cadets is also shown in the same plot (grey line, N=65). The vertical red dashed line is at the threshold value of 540, which is considered a differentiating point of physical fitness by the US Army. The horizontal black dashed line is at 0.5 or the median of the CDF. The male study cohort scored significantly higher than the larger USMA population (p=0.004, Kolmogorov-Smirnov test) **D.** Distribution of ACFT total points in the male study cohort, along with a vertical red dashed line at the 540 ACFT score cut-off; 60% of the male participants had scores above 540. **E.** Heatmap of the male cohort (rows) and their individual points for the six ACFT athletic events in the vertical columns (MDL, SPT, HRP, SDC, LTK, 2MR), where the darker grays indicate higher scores, and purple indicates a maximum score for that activity. **F.** CDF of the ACFT total points for Spring 2021 USMA West Point female total population (thick black line, N=493); the study cohort of female cadets is shown as the grey line (N=14). The vertical red dashed line is at the threshold value of 540 and the horizontal black dashed line is the median of the CDF. The female study cohort scored significantly higher than the larger USMA population (p=0.003, Kolmogorov-Smirnov test). **G.** Distribution of ACFT total points in the female study cohort, along with a vertical red dashed line at the 540 ACFT score cut-off. **H.** Heatmap of the female cohort (rows) and their individual points for the six ACFT athletic events in the vertical columns (MDL, SPT, HRP, SDC, LTK, 2MR), where the darker grays indicate higher scores, and purple indicates a maximum score for that activity.

During the three-month study, subjects participated in up to five separate study segments over which measurements were collected on body composition, cardiopulmonary performance (VO2max), personality and behavioral (NEO PI-3), and cognitive testing (Raven Progressive Matrices, Wisconsin Card Sorting Test (WCST)). Sleep and daily activity tracking measures were gathered throughout the study as well. Furthermore, within each segment, the subjects were exposed to bouts of intense cycling, ∼70% of their VO_2_ max, for a duration of 20 minutes. Cognitive testing was conducted pre- and post-exercise to evaluate the impact of physical stress. During each segment, four separate intravenous blood samples were collected: one at baseline prior to exercise, and three after exercise (approximately <5, 10, and 30 minutes, **Figure 1B**). For each study segment, cadets were randomly assigned to three periods of time throughout the day for their bike/blood draw events. This blood collection schedule is more representative of cadets’ everyday life and accounts for variations in the circadian rhythm. The blood samples were analyzed using an in-depth multi-omics approach which included untargeted metabolomics, untargeted proteomics, 3’mRNA sequencing, and EPIC array-based DNA methylation. In addition to these comprehensive multi-omics assays, targeted metabolic (iStat based) and cytokine panels, hematology and immunophenotyping analyses were performed at multiple blood draws. All molecular data were curated and annotated as described in the **Methods**. A total of 50,057 unique molecular measures were analyzed, including 16,544 gene regions and 16,415 gene promoters (DNA methylation), 16,318 mRNA transcripts, 139 untargeted proteins, 571 untargeted metabolites, 10 targeted metabolic biomarkers, 14 targeted cytokines, 38 hematological and 8 immunophenotypic measures. **Supplementary Table ST1** contains a detailed description of timepoints, subject numbers per segment and average number of molecular assays conducted per draw. After data curation and annotation, the final dataset used for analysis included 147 phenotypic features and ∼50,011 molecular analytes.

In **Figure 1C**, the empirical cumulative distribution functions (CDF) of ACFT total points are compared for the male study participants and the total male USMA population that took the ACFT in Spring of 2021. The study cohort (grey line in **Figure 1C**) scored significantly higher on ACFT than their counterparts in the USMA population (Kolmogorov-Smirnov test p-value = 0.004). The histogram (**Figure 1D**) presents the distribution of ACFT total points scored by the male cadets in this study, who had a median score of 553. An ACFT score of 540 has been proposed as a differentiating point of physical fitness by the US Army and male soldiers who score above this threshold have exemption from a body mass index assessment based on height and weight.^37^ Sixty percent of the male study participants attained an ACFT score >540. In this study, we used 540 (dotted vertical red lines in **Figures 1C** and **1D**) as the threshold score to differentiate higher- and lower-performers among the male participants. **Figure 1E** shows a heatmap of the six ACFT athletic events and the distribution of the points awarded across them (columns) for each of the male participants (rows). Four out of sixty-five male subjects achieved a perfect score of 600 on the ACFT. The heatmap also indicates that for some of the ACFT events, that there was a ceiling effect given that a significant portion of the subjects obtained the maximum score of 100 points for a single ACFT event.

The corresponding CDF, histogram and heatmap of female ACFT performance is presented in **Figures 1F, 1G,** and **1H** respectively. The ACFT measures were obtained on 14 of the 15 enrolled females. The average ACFT total points for the fourteen female subjects was 464 with all scores falling under the 540 ACFT threshold. However, similarly to the male subjects, the study female participants scored significantly higher on ACFT than their female counterparts in the USMA population (Kolmogorov-Smirnov test p-value = 0.003).

A summary of key phenotypic and demographic measurements broken out by gender is presented in **Table 1**. A more detailed list can be found in **Supplementary Table ST2** for phenotype features and **Supplementary Tables ST3-8** for iSTAT, metabolomic, proteomic, cytokine, transcriptomic, and DNA methylation assay measurements respectively. The NEO-PI personality tests results were not significantly different between the male and female groups. The activity tracking by the Garmin watches showed a trend of more cardio and strength activities for the female group although the male group tended to expend more average calories per activity. Sleep patterns were not significantly different as recorded by Oura rings. The men were significantly larger in height, body mass, skeletal muscle mass, and were able to generate more power as measured on the exercise bike.

**Table 1.** Summary of Demographic and Phenotypic Measurements. Mean (SD) reported for the male and female subjects along with the Bonferroni adjusted p-value from a two-sample Wilcoxon test between the two groups.

| Feature | Category | Male<br>(N = 71)<br>Mean (SD) | Female<br>(N = 15)<br>Mean (SD) | Bonferroni<br>adjusted<br>p-value |
| --- | --- | --- | --- | --- |
| Age at Enrollment (years) | Age | 20.7 (1.4) | 20.2 (0.7) | 1 |
| Body Height (meters) | Body Composition | 1.8 (0.1) | 1.6 (0.1) | 1E-05 |
| Body Mass (kg) | Body Composition | 81.5 (9.6) | 64.7 (7.4) | 2E-04 |
| BMI (kg/m<sup>2</sup>) | Body Composition | 25.0 (2.6) | 23.9 (1.4) | 1 |
| Body Fat (%) | Body Composition | 12.5 (4.4) | 22.0 (4.2) | 8.E-05 |
| Neuroticism (T-Score) | NEO-PI | 47.2 (10.7) | 52.2 (12.7) | 1 |
| Extraversion (T-Score) | NEO-PI | 52.6 (11.0) | 47.5 (8.5) | 1 |
| Openness (T-Score) | NEO-PI | 53.7 (10.8) | 56.4 (10.5) | 1 |
| Agreeableness (T-Score) | NEO-PI | 49.5 (11.7) | 50.7 (11.5) | 1 |
| Conscientiousness (T-Score) | NEO-PI | 58.6 (10.2) | 61.1 (8.1) | 1 |
| Correct Raven Progressive Matrix | Raven Progressive<br>Matrices | 14.0 (3.7) | 15.0 (2.6) | 1 |
| WCST Pre-Stress Total Errors Score | WCST | 18.4 (12.3) | 23.0 (22.0) | 1 |
| WCST Post-Stress Total Errors Score | WCST | 10.8 (5.1) | 12.3 (14.3) | 1 |
| Oura Sleep Time Avg (hours) | Oura Sleep | 6.0 (0.6) | 6.6 (0.9) | 1 |
| Oura Deep Sleep Avg (hours) | Oura Sleep | 1.9 (0.4) | 1.8 (0.5) | 1 |
| Oura REM Sleep Avg (hours) | Oura Sleep | 0.8 (0.3) | 1.1 (0.4) | 1 |
| Oura Resting Heart Rate Avg (bpm) | Oura Sleep | 47.1 (4.3) | 51.2 (5.7) | 1 |
| Oura HRV RMSSD/ms Avg | Oura Sleep | 88.8 (36.7) | 78.5 (32.0) | 1 |
| Number of Cardio Activities | Garmin Activity | 13.5 (11.0) | 24.0 (25.1) | 1 |
| Cardio Activity Avg Calories | Garmin Activity | 658.0 (475.0) | 375.3 (184.1) | 0.6 |
| Number of Strength Activities | Garmin Activity | 4.8 (8.9) | 13.8 (26.9) | 1 |
| Strength Activity Avg Calories | Garmin Activity | 294.3 (145.7) | 169.4 (70.4) | 1 |
| Average Power Bike Exercise Event 2 | Bike Metrics | 197.8 (62.4) | 143.1 (23.7) | 0.002 |
| Average Heart Rate Bike Exercise Event 2 | Bike Metrics | 156.8 (11.8) | 159.3 (16.7) | 1 |
| Army Combat Fitness Test | ACFT | 549.5 (37.1) | 463.9 (29.4) | 2E-05 |
| ACFT Maximum Deadlift (pounds) | ACFT | 303.4 (42.5) | 207.1 (32.9) | 1E-05 |
| ACFT Standing Power Throw (meters) | ACFT | 10.4 (1.7) | 6.2 (1.1) | 2E-06 |
| ACFT Hand Release Pushups (reps) | ACFT | 47.9 (9.6) | 42.5 (11.5) | 1 |
| ACFT Sprint Drag Carry (seconds) | ACFT | 94.1 (9.6) | 120.9 (11.8) | 2E-05 |
| ACFT Leg Tuck OR Plank (reps) | ACFT | 16.2 (5.0) | 9.3 (5.4) | 0.006 |
| ACFT 2 Mile Run (minutes) | ACFT | 13.9 (1.7) | 15.1 (1.3) | 0.057 |

### ACFT Associations with Phenotypic and Molecular Measurements

We explored which individual phenotypic measurements were associated with higher performance on the ACFT. These measurements included cardiopulmonary response to intense exercise and maximal oxygen uptake (VO_2_max), body composition (measured via bioelectrical impedance), personality (NEO-Personality Inventory (PI)), sleep (Oura ring), and activity tracking (Garmin watch). Spearman correlations were computed between the ACFT and its six individual event scores vs the phenotypic measurements for the entire cohort, only males, and only females with results reported in **Supplementary Table ST9**. The phenotype measures with the largest correlations to the ACFT Total Points were Skeletal Muscle Mass (r = 0.64) and the average power that a subject could generate during each of the four bike events (r = 0.54 to 0.68). These correlations were observed for both male and female subgroups, with slightly lower magnitude for the latter. VO2max was also found to have a statistically significant correlation (r = 0.41) with ACFT Total points as well as the specific ACFT event of 2-Mile Run for both males (r = -0.43) and females (r = -0.43). Resting Heart Rate as measured by the Oura Ring was on the edge of being significant with a Spearman correlation of -0.46 (Bonferroni adjusted p-value = 0.037).

We next explored the molecular omics measurements associated with ACFT. The omics types included in this analysis included metabolomics, proteomics, transcriptomics, DNA methylation as well as targeted cytokines and iSTAT measurements. Our first approach was the simplest in which the median value of a specific molecular analyte across all study segment events and blood draws was used to compute a correlation with the ACFT. The computed Spearman correlation for each if these omics is summarized in **Supplementary Tables ST10-15**. For iSTAT (**Supplementary Table ST10**), the top three analytes with the highest correlation to ACFT were urea (r = 0.59), creatinine (r = 0.52), and the protein creatine kinase (r = 0.4), all of which were statistically significant after adjusting for multiple comparisons (p-value < 0.006). For metabolomics (**Supplementary Table ST11**), there were twelve analytes that were statistically significant (Bonferroni adjusted p-value < 0.03), all positively correlated with ACFT Total Points (r = 0.46 to 0.63). Both urea and creatinine again appeared along with the vitamin C metabolite ascorbic acid 3-sulfate, several acylcarnitines (isovalerylcarnitine, isobutyryl-L-carnitine, and tiglylcarnitine), 2-Keto-glutaramic acid, 4-Deoxythreonic acid, and a diacylglycerol (DAG). For proteomics (**Supplementary Table ST12**), there was only one that was statistically significant, the complement component 4 binding protein (r = 0.46, Bonferroni corrected p-value = 0.014). There were no significant associations of the targeted Cytokines (**Supplementary Table ST13**) with ACFT Total points. Finally, for transcriptomics (**Supplementary Table ST14**) of whole blood, there were two gene transcripts that were correlated in a statistically significant way with ACFT Total points: gamma-secretase activating protein (GSAP, r = 0.55, adjusted p-Value = 0.007) and the long non-coding JPX transcript, XIST activator (JPX, r = -0.54, adjusted p-Value = 0.011). JPX is known to upregulate the RNA expression of XIST. Both the JPX and XIST transcripts are significantly higher in women vs. men (adjusted p-value 0.029 and 3E-05 respectively from **Supplementary Table ST7**).

Our second approach to analyzing the association of the molecular measurements with ACFT considered the longitudinal dynamics of the study segments occurring over multiple weeks and the temporal dynamics of individual blood draws collected pre- and post-exercise (70% VO2max cycling). We conducted mixed-effect ANOVAs that used the ACFT Total Points grouping defined by the threshold of 540 (**Figure 1C**) as the “between subject factor”, whereas the “within subject factors” were defined as the study segment and blood draw for one analysis. The mixed-effect ANOVA analysis was only applied to the male subjects for two reasons: 1-the threshold of 540 is specified for the male population and there is no equivalent Army defined threshold for females; 2-the number of female subjects in our study was considerably lower than the males. We also conducted an alternative analysis where we replaced the blood draw factor with fold change measures defined as blood draw two (<5 minutes), three (∼10 minutes) and four (∼30 minutes post bike exercise) divided by the blood draw one measure that occurred just before the individual started riding the bike for 20 minutes at 70% VO2max. It should be noted that within each study segment, there were many analytes that changed significantly across the four blood draws in response to intense physical exertion. However, the overall dynamic response to exercise was similar across the low and high performing sub-groups. The results from the mixed-effect ANOVAs in male subjects can be found in **Supplementary Tables ST16-27,** where many of same analytes that were identified to be significantly correlative to ACFT from the median Spearman correlation analysis (e.g., urea, the acylcarnitines, 2-keto-glutaramic acid, complement component 4 binding protein, etc.) were also found to be significantly correlated with the groupings of the ACFT performance by the 540-threshold (See representative examples in **Supplementary Figure S1**).

### Identifying Expression Circuits Associated with ACFT Performance

We next wanted to investigate how groups of features (i.e., small molecules, proteins, transcripts) that interact with one another might be associated and how these groups could be predictive of ACFT. Therefore, we developed an analytics pipeline that we call PhenoMol, which aims to identify subsets of molecular features that are associated with a given outcome metric through a network approach. PhenoMol reconstructs biological networks using multi-omics measurements and an interactome of known molecular interactions. The interactome is a graph built from public repositories of protein-protein and protein-metabolite interactions.^29,30^ and represents all known interactions between molecules involved in human biological processes at the time of analysis. Molecular subnetworks associated with a phenotype or outcome are generated by solving the prize-collecting Steiner forest (PCSF) algorithm on the interactome to find subnetworks that collect node prizes and reduce edge costs. An example of a node prize is the absolute Spearman’s correlation between the molecular features represented by a node on the network and the outcome performance metric of interest. An example of an edge cost between two nodes is proportional to one minus the fractional confidence level of the interaction between two nodes. PhenoMol’s application of PCSF network graph-based method is based on the open access software tool called Omics Integrator (OI).^31^

**Figure 2** depicts an overview of the method used to generate PhenoMol models compared to the analogous statistical approach without the PCSF dimensionality reduction step (control models). Cross-fold stratified subsampling of the cohort is utilized to limit overfitting for both PhenoMol and the control models (**Figure 2A**). The control models are trained using Sparse Partial Least Squares Regression (sPLSR) on 80% of the data and tested on the remaining 20% of data. The input molecular measures for both PhenoMol and the control models were the computed median values across the different blood draw times and study segments (**Supplementary ST1**). The choice of median molecular measures was driven by two factors. Due to the challenges of field research in a military operational environment, during data collection from active-duty USMA cadets (including safety and time constraints due to on-going training and class schedules) some traditional study practices, such as circadian-matched replicates, fasting blood draws and regular sleep schedules, could not be implemented. The median molecular data serve both to limit the influence of any individual testing event as well as to approximate the variability of subject condition likely to be found when implementing assessment techniques like this in field operations. To prevent the confounding of effect of gender on both the phenotype and molecular measures, the analysis was run using only the 65 male subjects.

**Figure 2.**
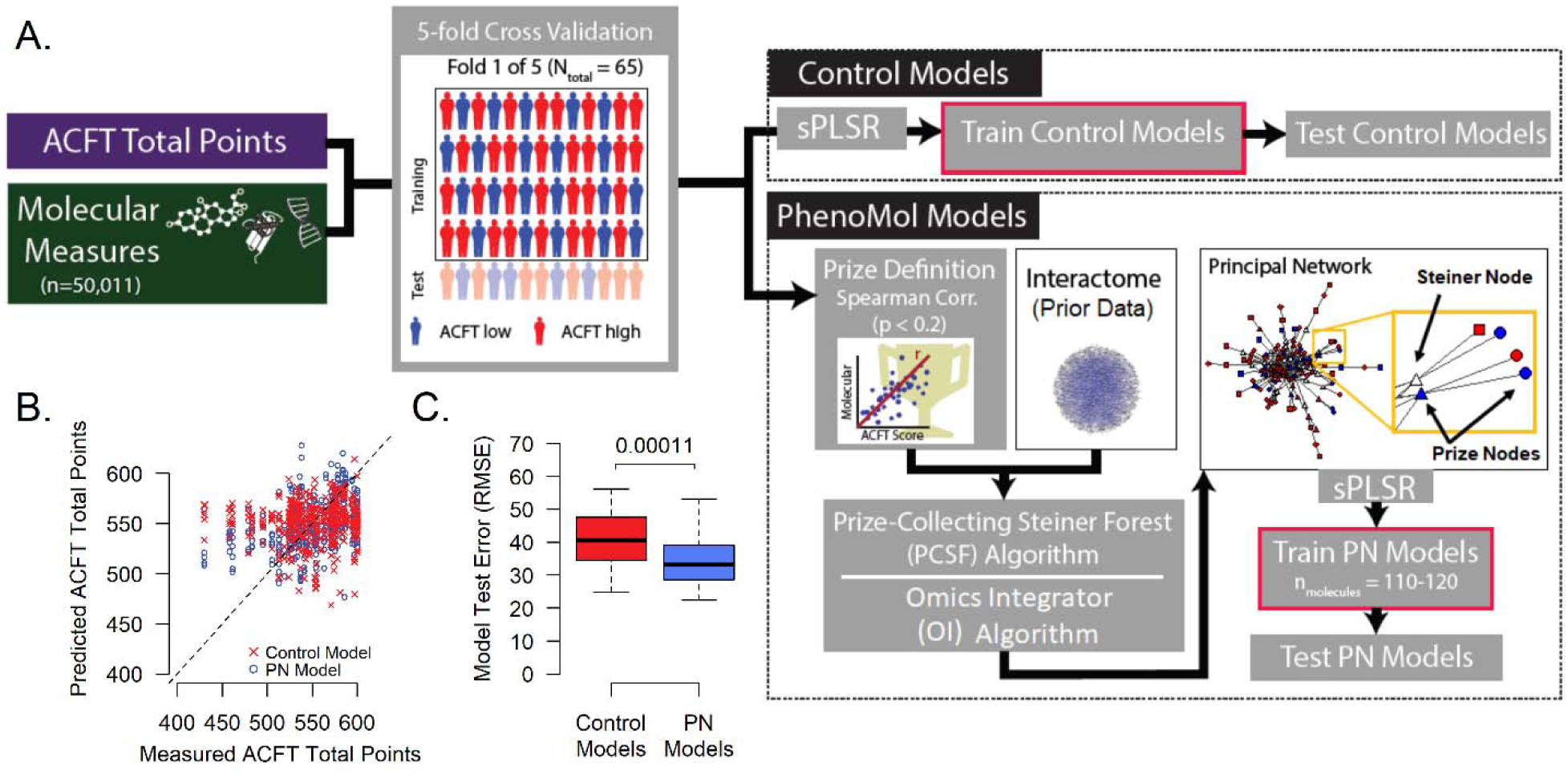
Unsupervised Reconstruction of Biological Networks Associated with Army Combat Fitness Test Performance: **A.** Overview of the method used to generate the PhenoMol Models in comparison to a simpler modeling approach (i.e., Control Models). In both cases, a 5-fold stratification subsampling of the cohort was used to control overfitting. The control models are trained using Sparse Partial Least Squares Regression (sPLSR) on 80% of the data and tested on the remaining 20% of data. The PhenoMol generated models includes a network-based data reduction methodology that utilizes the Prize Collecting Steiner Forest (PCSF) algorithm and the Omics Integrator (OI) algorithm. The inputs to these algorithms include both prizes and an interactome of prior data. The output of this analysis is the Principal Network (PN), which contains a set of prize nodes that were measured and Steiner nodes that were not measured but were robustly selected by the PCSF algorithm. A PN model is then generated by sPLSR that only utilizes molecular features found within the PN. **B.** Scatter plot presenting the predicted vs measured ACFT Total Points for the 65 male cadets for the PN models (blue circles) and the control models (red x’s) for sPLSR generated models across six analysis runs of five-fold cross validations (N = 30). **C.** Box plot presenting the root mean square error (RMSE) for predicting the ACFT Total Points of 65 male cadets. The box plot presents the RMSE for thirty sPLSR models (i.e., six analysis runs of five-fold cross validations). The RMSE for the PN models was significantly lower than the control models with a two-sample paired Wilcoxon signed rank test p-value = 0.00011.

The PhenoMol models are also generated using sPLSR, trained and tested in the same cross-fold validation process as the control models. For this analysis, the contraints imposed by the network-based data reduction methodology that utilize PCSF algorithm were different. The input to these algorithms includes both prizes and an interactome of prior data. The prizes are the set of absolute Spearman’s correlations between the ACFT Total Points and each measured molecular feature for the cohort that meet a threshold p-value < 0.2. The interactome of prior data includes all known molecular interactions. The PCSF algorithm is used by the OI algorithm to generate multiple subnetworks (N = 100) in which noise is added to the edge costs. The robustness of a node is then computed based on its frequency of appearance in all of the subnetworks. Nodes achieving a robustness greater than 20% of the subnetworks are retained. These retained nodes and all known interactions between the retained nodes form a graph, which we label the Principal Network (PN). The PN contains two types of nodes: nodes with an associated prize, which correspond to molecules in the measured dataset (terminal nodes) and nodes without an associated prize, which correspond to molecules connected to terminal nodes by graph edges (Steiner nodes). A PN model is then generated by sPLSR that only utilizes molecular features found within the PN. The PNs that were generated across the folds contained 110 to 120 prize nodes (measured molecular features) which is a significant reduction from the original 50,011 measured molecular features. Scatter plots presenting the predicted vs. measured ACFT Total Points for the 65 male cadets for the PN models (blue circles) and the control models (red x’s) are shown in **Figure 2B**. A box plot representing the root mean square error (RMSE) of ACFT Total Points prediction for the 65 male cadets are shown in **Figure 2C**. The RMSE for the PN models was significantly lower than the control models with a two-sample paired Wilcoxon signed rank test p-value = 0.00011.

After demonstrating that the data-reduced PNs performed better than the control models, we next applied Louvain clustering to the PN network to identify modules of tightly connected molecules (See **Figure 3A** and **Figure 3B**), referred to as Principal Network Modules (PNMs). For the PN generated in each fold of cross-validation, Louvain clustering generated up to seven PNMs. The PNMs were labeled numerically in order of their size, i.e., the number of nodes in the PNM. The location of the PNMs on the grid (Figure 3B) is such that the number of common edge connections of each PNM and its immediate neighboring clusters are maximized for visualization purposes. A total of 205 PNMs were generated from six analysis runs of five-fold cross validations (N = 30), with an average of ∼7 PNMs per fold. **Figure 3C** shows the overlap of the PN nodes across the 5-folds, while the heatmap in **Figure 3D** presents the agglomerative hierarchical clustering (Ward Method) of the fraction of node overlap between 205 PNMs. The heatmap demonstrates that the PNMs being generated from the PN within each fold of cross-validation represent similar sets of common nodes. The emergence of PNMs with common nodes across the folds indicates robust network substructures that can be evaluated further. We subsequently generated PNM based models to predict ACFT performance by applying sPLSR with the outcome (ACFT Total Points) as a target function and limiting the molecular features to those that were members of the PNM. Ensemble PNM models were then generated for all combinations using a N-Choose-K process where K ranged from 1 to N when choosing from the N PNMs. In Figure 3F, the RMSE of the PhenoMol models (including the PN models, the PNM base models and ensemble PNM models) and control models are compared. As expected, the RMSE of individual PNM base models that include a smaller set of nodes do not perform as well as the ensemble PNM models. Because the PNMs represent and encode subsets of tightly interacting molecular nodes within a PN, the PNM ensemble models provide the means to annotate and biologically interpret the resulting predictive models more than is possible with the PN model, while still achieving comparable predictive performance.

**Figure 3.**
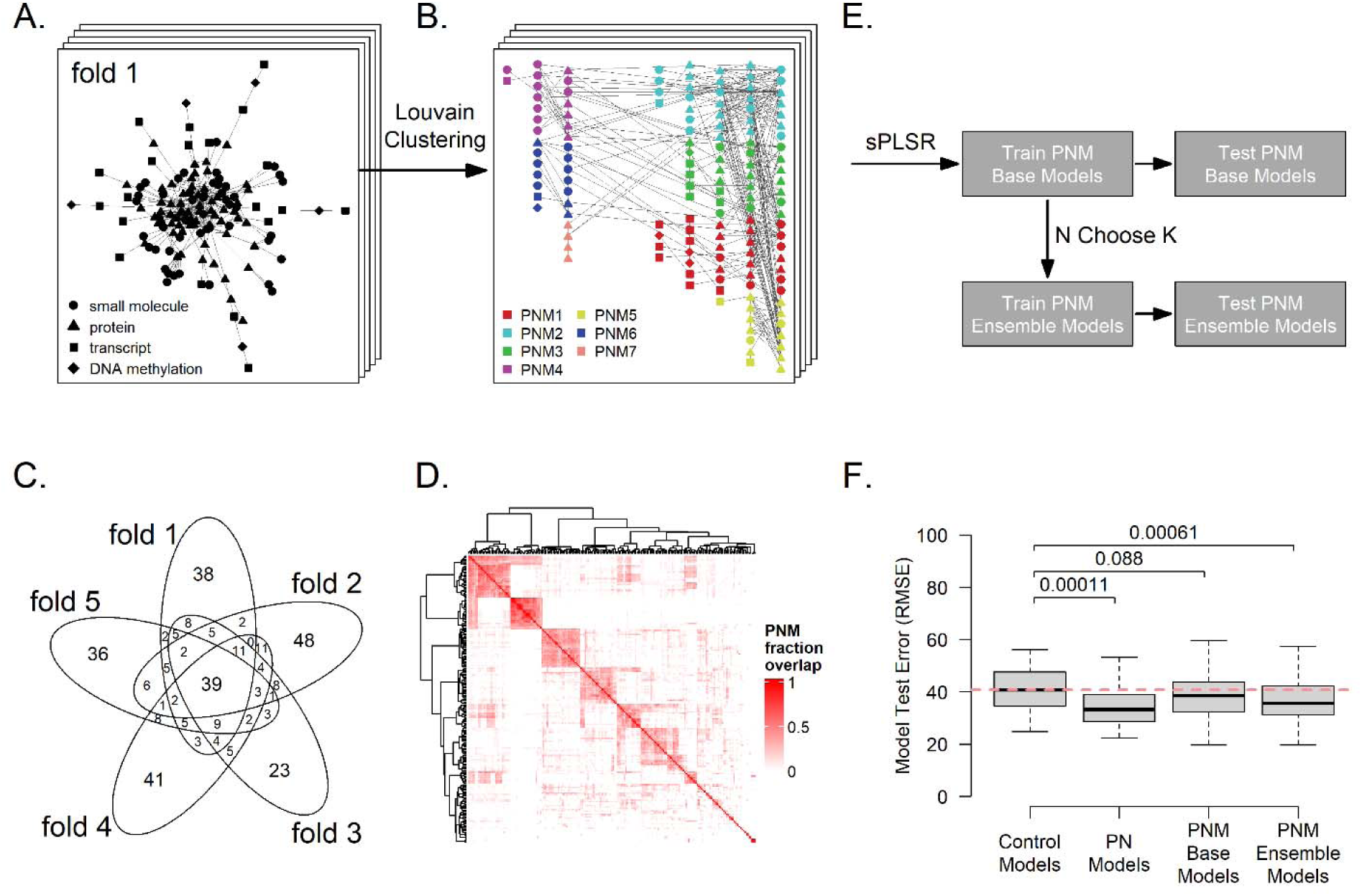
Identification of Principal Network Modules (PNMs): **A.** Louvain clustering is applied to the Principal Network (PN) to decompose it into Principal Network Modules (PNMs) within each fold. **B**. A total of seven PNMs were generated from the PN of fold 1 and are labeled numerically in order of their size (i.e. number of nodes). The nodes of the PN have been color coded by their PNM membership and replotted in a grid layout. **C.** The Venn diagram presents the overlap of the PN nodes across the 5-folds. There were 39 (28%) of the ∼ 139 nodes per fold (average) present in all five folds and 37.2 (27%) unique nodes observed in each of the five folds. **D.** The heatmap presents the agglomerative hierarchical clustering (Ward Method) of the fraction of node overlap between 205 PNMs. The heatmap color scaling ranging from 0% (white) to 100% (red) overlap of nodes between pairs of PNMs. These 205 PNMs were generated from six analysis runs of five-fold cross validations (N = 30) that upon clustering led to 6.83 PNMs per fold on average. **E.** PNM base models were generated and trained by applying sPLSR with respect to the outcome of ACFT Total Points and limiting the molecular features to those that were members of the PNM. Ensembles of the PNM models were then generated for all combinations using a N choose K process where K ranged from 1 to N when choosing from the N PNMs. **F.** The box plot presents the RMSE when testing the control models, PN models, the PNM base models, and the PNM Ensemble models. The model test error was compared between model types with the reported p-values reported generated from a two-sample paired Wilcoxon signed rank test. The red horizontal dashed line is at the median RMSE for the control models.

We next investigated methodologies to integrate composite performance metrics from multiple outcome scores. As mentioned earlier, ACFT is a composite score on a scale up to 600 points and aggregates six separate athletic events (**Figure 1A**), each contributing a maximum of 100 points to the total score. It is straightforward to model each ACFT event separately with corresponding output models from PhenoMol. This would lead to PNs and PNMs that are trained and specific to each individual ACFT event. However, the ACFT events are not entirely independent of each other. For example, the sprint drag carry includes running as does the 2-mile run event. The same biology that is relevant to one can also be important to the other. At the same time, both events have other characteristics that are distinct from each other. For example, the sprint drag carry event includes lifting, carrying, and dragging heavy weights, unlike the 2-mile run. PhenoMol was developed so that it can integrate multiple sub-networks generated from multiple outcome performance metrics into a single PN. This integration of multiple sub-networks was achieved by taking the union of the nodes from the set of sub-networks (**Figure 4A, 4B**) and completing it on the interactome. As described previously, Louvain clustering is applied to separate the PN into Principal Network Modules (PNMs) (**Figure 4C**). The union of the subnetworks from the individual ACFT events leads to a richer PN and PNMs than the PN and PNMs associated with the total ACFT Total Points, as demonstrated by comparing the PNs (**Figure 4B vs Figure 3A**) and PNMs (**Figure 4C vs Figure 3B**). The reason for this increased richness is due to modeling of granular sub-scores and integration of more fundamental contributors to the outcome of ACFT Total Points.

**Figure 4.**
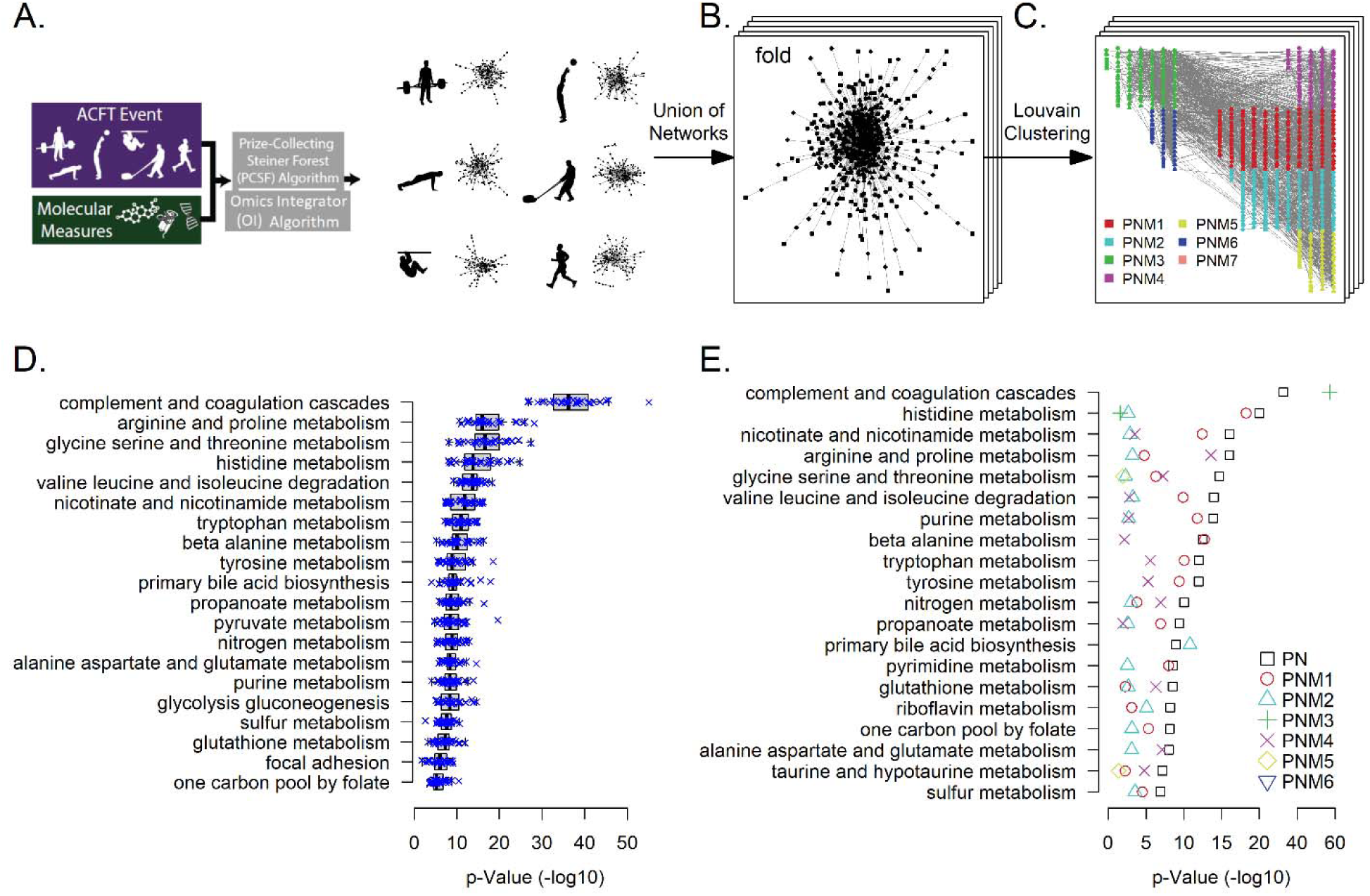
Biological Expression Circuits of the Army Combat Fitness Test: **A.** Generated a set of phenotype subnetworks, one for each of the six ACFT events that include Maximum Deadlift, Standing Power Throw, Hand Release Pushups, Sprint Drag Carry, Leg Tuck, and 2 Mile Run. **B**. The principal network (PN) is generated by taking the union of nodes from the set of six phenotype subnetworks. This is repeated for each fold of the data. **C.** Louvain clustering is then applied to the Principal Network (PN) to decompose it into Principal Network Modules (PNMs) within each fold. **D.** Pathway enrichment analysis was performed on the thirty PNs generated from the set of six analysis runs of five-fold cross validations (N = 30). The top twenty pathways are presented for each of the thirty PNs and ordered by the average p-value for each pathway. **E.** The top 20 pathways from enrichment analysis performed on the PN and the six PNMs from the exemplary fold (B and C) are shown. The ordering of the pathways is by the p-value of the enrichment analysis of the PN (union of the six PNMs). For each PNM, the pathway is considered enriched in the PNM if its p-value < 0.05.

Subsequently, we performed over-representation analysis (ORA) on the thirty PNs generated from the set of six analysis runs of five-fold cross validations (N = 30). This analysis allowed us to determine if the nodes within the PN are overrepresented in known biological pathways (**Figure 4D**). ^32^A benefit of this approach is that it does not require measurements for each node in the networks (as mentioned above, the PN generated by the PCSF algorithm includes Steiner nodes). In **Figure 4D**, the top twenty pathways are presented and ordered by the average enrichment p-value across the thirty PNs for each pathway. The complement and coagulation cascades pathway was identified in all thirty PNs and the most significant relative to the other pathways. The second top pathway identified was arginine and proline metabolism that includes the urea cycle. Enrichment analysis was next conducted on the set of PNMs generated from each PN. Figure 4E presents the top 20 pathways on the six PNMs from the exemplar fold (shown in **Figure 4C**). The ordering for the pathways in **Figure 4E** is by the enrichment p-value of the exemplar fold and once again the complement and coagulation cascades pathway is at the top. From the symbols on the figure one can determine which PNMs are enriched for each pathway (i.e., those that reach the p-value threshold value of 0.05). Note that PNM3 (green plus sign symbol) in **Figure 4E** contains a significant number of nodes that belong the complement and coagulation cascades pathway, while PNM4, and to a lesser extent PNM1 and PNM2 contain nodes that are enriched in arginine and proline metabolism. This is an example where the annotation of the PNMs (represent sets of tightly connected nodes within the PN) along with the edge structure that interconnects the two PNMs (PNM3, top left, and PNM4, top right of Figure 4C) can provide a rich contextual biological understanding when interpreting the resulting mathematical models (i.e., PNM ensemble models).

### Models for ACFT Scores Prediction

The PN and PNMs from the exemplar fold presented in **Figure 4** were used to generate Base and Ensemble models and trained using sPLSR as described above (**Figure 3E**). Six sets of models were trained to predict each of the six ACFT events that include Maximum Deadlift, Standing Power Throw, Hand Release Pushups, Sprint Drag Carry, Leg Tuck, and 2 Mile Run. **Figure 5B** presents the predicted (y-axis) vs. the measured (x-axis) ACFT events for the 65 male West Point cadets, where control models (red x’s) and the PNM Ensemble models (blue circles) are compared. For clarity purposes, the illustrative data presented here is from one fold of one analysis run. Overall, the predicted and measured ACFT scores are in good agreement, particulary for higher performing subjects. However, it should be noted that there is a ceiling effect for both the Maximum Deadlift and Leg Tuck. The test scoring and administration limited the cadets to lifting a maximum of 340 pounds and 20 reps for Leg Tucks. Publically available ACFT tables were used to convert a raw ACFT measurement from units of pounds, meters, minutes, etc. to ACFT event scores (scale of zero to one hundred (**Figure 5C**)). The ACFT Total Points is the summation of all six ACFT event scores and thus it ranges from zero to a maxium of 600 points. In **Figure 5D**, the PhenoMol PNM ensemble models (blue circles) and control models (red x’s) predictions versus measured ACFT Total Points for the 65 male West Point cadets from one fold of one analysis run are shown. The cross fold validation of the examplar ACFT Total Points model achieved a training R^2^ (mean ± standard deviation) of 0.687 ± 0.036 and a test R^2^ of 0.589 ± 0.068. Permutation testing was conducted in which the the subject ID labels for the molecular measures was randomly shuffled followed by running the entire analysis. This was repeated three hundred times to generate a distribution of R^2^ reflective of random occurances. Only one of the three-hundred permutations (p-value = 0.0033) achieved an R^2^ higher than 0.589. A box plot presenting the RMSE of ACFT Total Points prediction from a set of six analysis runs of five-fold cross validations (N = 30) are shown in **Figure 5E**. The RMSE for the PNM ensemble models was significantly lower than the control models with a two-sample paired Wilcoxon signed rank test p-value = 1E-7.

**Figure 5.**
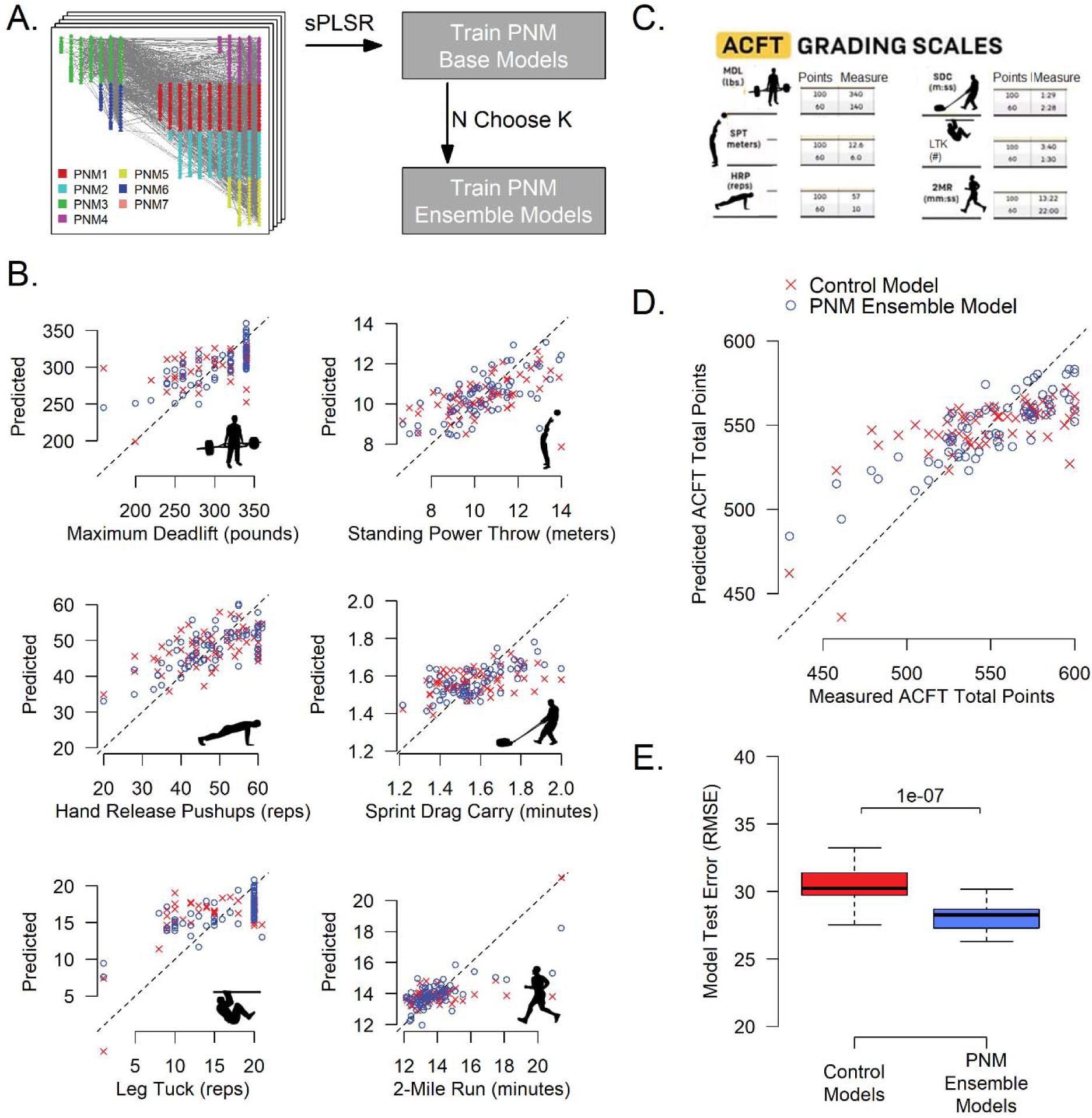
Machine Learning Models for Predicting Phenotypes of the Army Combat Fitness Test. **A.** PNM Base and Ensemble models were trained for each of the six ACFT events that include Maximum Deadlift, Standing Power Throw, Hand Release Pushups, Sprint Drag Carry, Leg Tuck, and 2 Mile Run. **B.** The predicted (y-axis) vs the measured (x-axis) are presented for the control models (red X’s) and the PNM Ensemble models (blue circles) for each of the six ACFT events for the 65 male West Point cadets. The illustrative data presented is from one-fold of one analysis run. **C.** The ACFT grading scale converts each of the ACFT event measures into a score that ranges from zero to one hundred points. The ACFT Total Points is the summation of all six ACFT event scores and thus it ranges from zero to a maximum of 600 points. **D.** The predicted vs. measured ACFT Total Points for the 65 male West Point cadets is shown for one-fold of one analysis run. The predicted data is from both the control models (red X’s) and from the PNM ensemble models (blue circles). **E.** A box plot presenting the RMSE for predicting the ACFT Total Points of 65 male cadets generated from a set of six analysis runs of five-fold cross validations (N = 30) is shown. The RMSE for the PNM ensemble models was significantly lower than the control models with a two-sample paired Wilcoxon signed rank test p-value = 1E-7.

To better characterize the PNM Ensemble Model that achieved the highest performance during testing, we generated a comprehensive summary which is presented in **Figure 6**. Starting first with the PNMs, **Figure 6A** shows the top biological annotations for each of the PNMs used to predict the ACFT events and total scores. In **Figure 6B**, the set of molecular features selected by sPLSR to model the six ACFT events are shown. Agglomerative hierarchical clustering (Ward Method) was performed on these molecular features. The columns in the heatmap represent the 65 male West Point cadets, where the subjects with a high measured ACFT Total Points > 540 (red) or < 540 (blue) are highlighted in the horizontal bar at the top of the heatmap. The omic type and the PNM for each of the molecular features are also indicated by the left side vertical bars. A network view of the molecular features from the heatmap along with the shortest distance connecting edges is shown in **Figure 6C**. The PNM contributions to each ACFT event model is illustrated in **Figure 6D**, where the percent contribution is computed from the PNM’s molecular feature summed square model loadings. The top biological annotations for PNM3 and PNM2, which are the largest contributors to the PhenoMol models of ACFT, are the complement and coagulation casade and primary bile acid biosynthesis, respectively.

**Figure 6.**
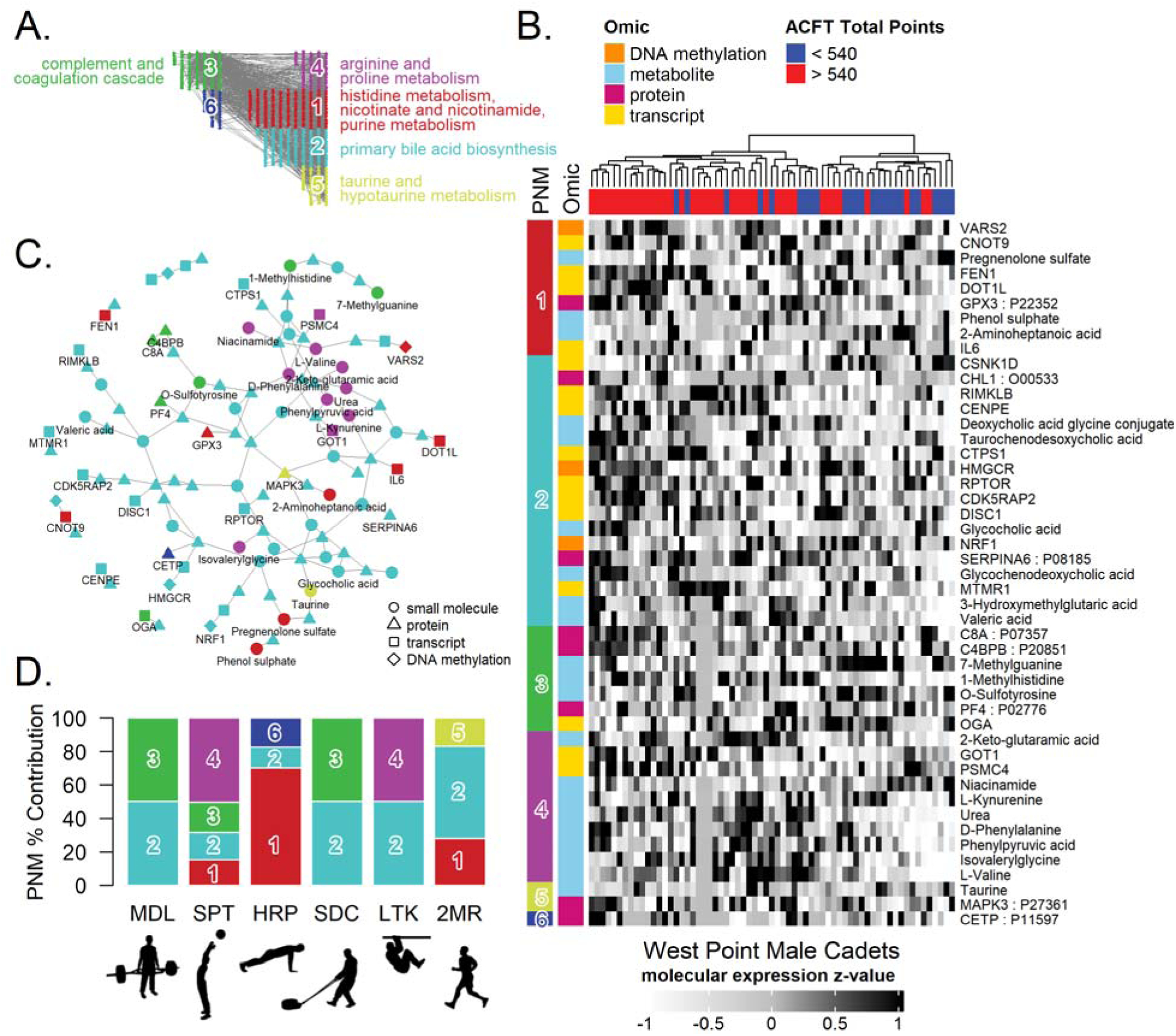
Integrated network-based analysis of multiple phenotypes and omics measurements: **A**. Top biological annotations for each of the PNMs. **B.** Heatmap of expression values for the set of molecular features selected by sPLSR to model the six ACFT events. The columns represent the 65 male West Point cadets and have undergone agglomerative hierarchical clustering (Ward Method). Those subjects with a high measured ACFT Total Points > 540 (red) or < 540 (blue) are highlighted in the horizontal bar at the top of the heatmap. The omic type and the PNM for each of the molecular features is indicated by the left side vertical bars. **C**. Network view of the molecular features from the heatmap along with the shortest distance connecting edges. **D.** PNM contributions to each ACFT event model. The PNM percent contribution is computed from the PNM’s molecular feature summed square model loadings.

### Surrogate Phenotype Features of the Army Combat Fitness Test

Phenotypic models were generated by sPLSR for each of the six ACFT events from 140 phenotypic measures on the 65 male West Point cadets across six analysis runs of five-fold cross validations (N = 30). The features selected by sPLSR in 50% or more of these 30 folds are grouped in four categories and shown in **Figure 7A** and described in more detail in **Supplementary Table ST28**. **Figure 7B** compares the predicted vs. measured ACFT events using the Phenotypic Models (black X’s) and omics-based PNM Ensemble models (blue circles). As shown in this figure, the performance of phenotype- and omics-based models are comparable. A closer inspection of the results, highlights a ceiling effect for both the Maximum Deadlift and Leg Tuck events. The test scoring and administration limited the cadets to lifting a maximum of 340 pounds and 20 reps for Leg Tucks. The pie charts in **Figure 7B** present the contributions to the phenotypic models by feature class. **Figure 7C** shows the predicted vs. measured ACFT Total Points for the from one fold of one analysis run, where the Phenotypic models (black X’s) and the omics-based PNM Ensemble models (blue circles) are included. The root mean square error (RMSE) of predicting the ACFT Total for the phenotypic models is slightly lower in test error than the omics based PNM ensemble models with a two-sample paired Wilcoxon signed rank test p-value = 2.6E-4.

**Figure 7.**
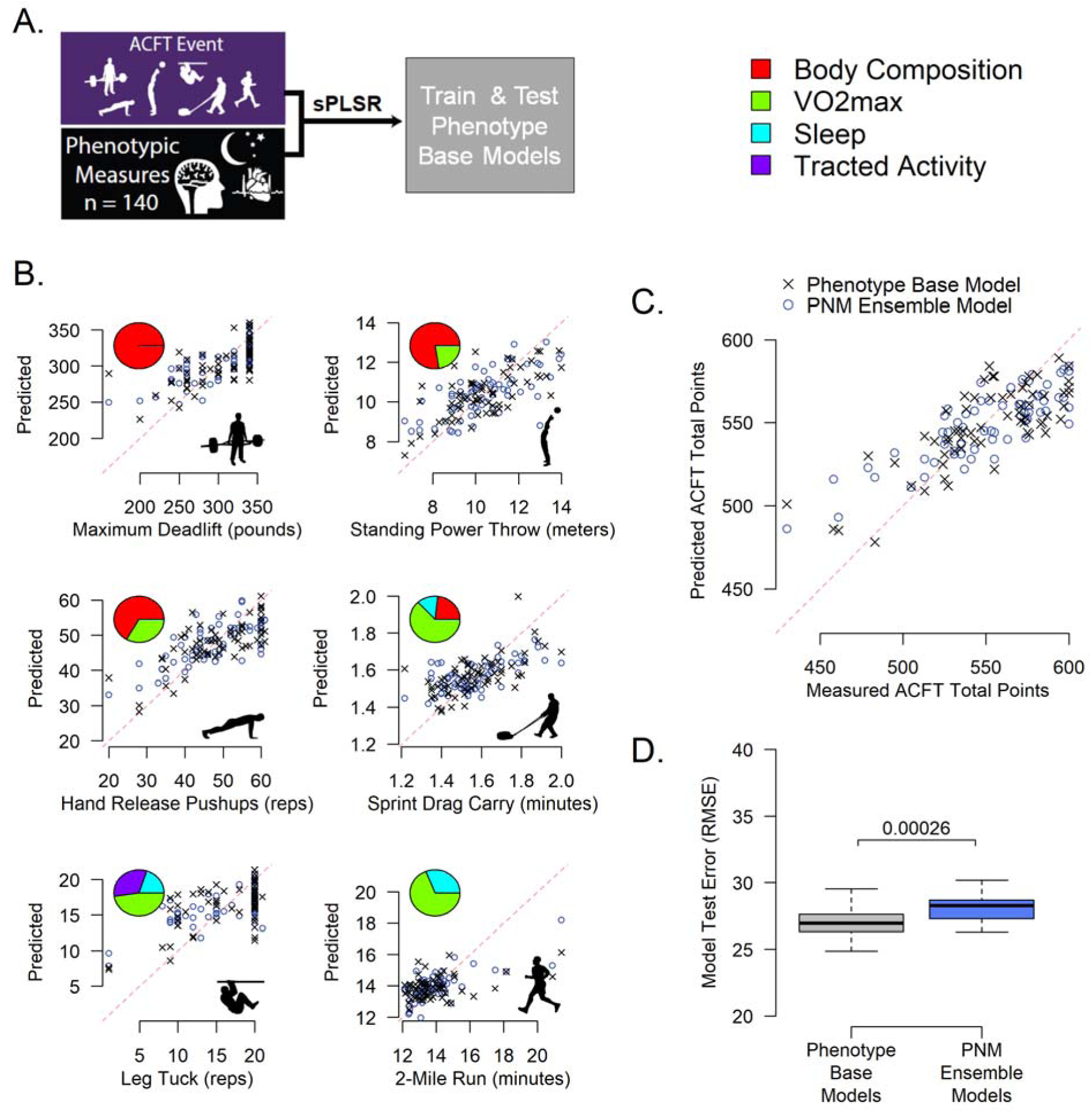
Surrogate Phenotype Features of the Army Combat Fitness Test. **A**. Phenotypic models were generated by sPLSR for each of the six ACFT events from 140 phenotypic measures on the West Point male cadets. Models were generated for each ACFT event across six analysis runs of five-fold cross validations (N = 30). **B.** The predicted (y-axis) vs the measured (x-axis) are presented for the Phenotypic models (black X’s) and the omics-based PNM Ensemble models (blue circles) for each of the six ACFT events for the 65 male West Point cadets. The illustrative data presented is from one-fold of one analysis run. There is a ceiling effect for both the Maximum Deadlift and Leg Tuck. The test scoring and administration limited the cadets to lifting a maximum of 340 pounds and 20 reps for Leg Tucks. The pie charts present the contributions to the phenotypic models by feature class for each of the six ACFT events. The contributions shown are the top classes of features that were selected by sPLSR in 50% or more of these 30 folds for each ACFT event. **C.** The predicted vs measured ACFT Total Points for the 65 male West Point cadets from one-fold of one analysis run. The predicted data is from both the Phenotypic models (black X’s) and the omics-based PNM Ensemble models (blue circles). **D.** Box plot presenting the root mean square error (RMSE) for predicting the ACFT Total Points of 65 male cadets generated from a set of six analysis runs of five-fold cross validations (N = 30). The RMSE for the phenotypic models are slightly lower in test error than the omics-based PNM ensemble models with a two-sample paired Wilcoxon signed rank test p-value = 2.6E-4.

## DISCUSSION

The rapid development and diversification of technologies for molecular measurement create an opportunity to identify molecular features associated with complex traits of human performance. One complicating factor is the relatively small cohort sizes of most human subject studies, which limits applicability of purely data-driven statistical methods to discover significant associations among tens of thousands of analytes. To address this issue, we have developed a computational approach (PhenoMol) to tackle feature-rich behavioral, cognitive and molecular datasets collected from cohorts where the number of data points vastly outnumbers the cohort size. There are a variety of algorithms that exist for the integration of multi-omics data to identify specific groups, types, or states of either individual cells or subjects.^9,33^ These algorithms aim at integrating different molecular layers that include epigenetics, transcriptomics, proteomics, and metabolomics and relate them to the observed phenotypes of health and disease status at the individual level. Many of the computational approaches that have been explored are linear algorithms, which are often more robust due to small numbers of tuning parameters.^9,33^ However, these methods can lack in accuracy by not accounting for non-linearities that regularly occur in complex biological systems. Neural networks have been applied to overcome these limitations^34^ but require large datasets. PhenoMol combines both linear (sPLSR) and non-linear (PCSF, network graph theory, Louvain clustering) methods and utilizes prior data on human biology (molecular interactome) to handle datasets with small numbers of subjects. Furthermore, many existing multi-omic integration approaches are either focused exclusively on pathway-level analysis of cellular and molecular biology data^33,34^ or only link phenotypic information to the underlying biology as a post-hoc analytic step.^9^ We developed a methodology that is explicitly built to 1) integrate all information from phenotypic data that is associated with outcomes of interest, 2) frame multi-omic integration as a non-linear graph optimization problem to discover biological networks that are coupled to observed phenotypic data, and 3) incorporate powerful constraints from prior knowledge of molecular interactions observed in human biology to reduce dimensionality of the problem and provide robustness in the face of modest sample sizes frequently encountered in studies of human performance.

PhenoMol was used to predict elite-level physical performance as defined by the U.S. Army Combat Fitness Test (ACFT). ACFT is a composite score and consists of six separate athletic events. Using the PhenoMol pipeline, we integrated the molecular sub-networks from each of these athletic events into a rich principal network to predict the outcome of interest (ACFT Total Points). This approach allowed us to capture the biological networks associated with the phenotypes of the outcome specifically. Surrogate phenotypes could also be used to predict the outcomes of interest, as shown in **Figure 7**. In this work, and consistent with prior literature reports, the ACFT sub-score of Maximum Deadlift, Standing Power Throw and Hand Release were predominantly described by the Body Composition surrogate phenotypic feature class, where individuals with more lean mass are expected to outperform at these events. ^35,36^ Similarly, VO2 max was the dominant feature class of phenotypic models for the Sprint Drag Carry and 2-Mile Run events. A side by side comparsion of the phenotypic and PhenoMol-based molecular models in **Figure 7** shows comparable outcome prediction capabilities for the male subjects. It should be noted that the predicted and measured ACFT scores are in good agreement, particulary for higher performing subjects; however, all models developed in this work were more limited in predicting the subjects with the lowest ACFT event measurements (for example see 2 mile run in **Figure 5B**). A plausible explanation for this observation is the makeup of our cohort. The recruiting pool for this study focused on USMA cadets who were undergoing extensive training and physical conditioning in preparation for the 2021 Sandhurst Military Skills Competition; an annual event at West Point comprised of US and international teams.^37^ As a result, more than sixy percent of the partcipating male subjects achieved ACFT Total Points higher than the US Army threshold of 540, which is significantly above the average ACFT performance of the USMA population (**Figure 1C**). Furthemore, the male study cohort included fewer low performing cadets (with ACFT scores below 510 points) compared to those who excelled at this physical fitness test. Therefore, this imbalance in the cohort can cause a bias of the models towards higher performing cadets.

Application of the PhenoMol analytics pipeline to the data collected in the USMA female cohort was determined not to be feasible due to the small number of female subjects enrolled in the study, which precludes the implementation of key steps of the analytics pipeline such as multi-fold cross training and validation in the female only dataset. The strength of the PhenoMol approach is that the constraints imposed by the network-based (PCSF and OI algorithms) can reduce fifty thousand measured molecular features down to a couple hundred when generating the principal network and further reduced this by Louvain clustering when generating the principal network modules, allowing the analysis of 65 male subjects. For the fourteen female subjects, which require splitting into training and test groups, the limitation of too many features in a PNM vs. the number of subjects becomes an issue. Moreover, the ACFT event and total scores are significantly different between the female and male populations (**Figure 1F-H**). Comprehensive studies have clearly shown the gender bias of the 2019 assessment and scoring system used in this study.^27^ Our observations are consistent with these prior studies, and as mentioned earlier the Army threshold value of 540 for males is not applicable to females. Similarly, we refrained from applying the analytics pipeline to the combined male and female dataset due to the significant imbalance of the cohort (65 male vs 14 female subjects with ACFT measures), statistical differences of the event scores, and confounding effects caused by the gender specificity of some analytes. It should be noted that ratio of male to females in this study is representative of the USMA demographics and our recruiting approach was gender agnostic. A comparative analysis of male versus female biological aptitudes for ACFT performance will require a larger study with equal representation of both sexes and is outside the scope of this work. However, we believe it is still important to report on our findings due to the scarcity of published data in elite female performers, including military personnel and athletes.

A key aspect of the PhenoMol pipeline is the incorporation of powerful constraints from prior knowledge of molecular interactions observed in human biology to reduce the dimensionality of the molecular data for subsequent outcome prediction. To this end, we built a graph-theory based method to identify the molecular mechanisms driving phenotypes of performance and organized it in expression circuits with identified biological functions. We define an expression circuit to have the following three characteristics: 1) must contain a relatively small set of measurable molecular features; 2) the set of measurable molecular features must be predictive of the phenotypes and/or outcomes of interest; 3) the set of molecular features must be linked mechanistically to one another. We use OI, a mechanistic model that integrates prior biological knowledge (human interactome) and integrates many omics streams including epigenome, genome, transcriptome, proteome, and metabolome, to distinguish functionally related molecules (true circuits) from purely statistical associations. Subsequent application of Louvain clustering allows us to uncover principal network modules (PNM) of tightly connected molecules that are also exhibit coordinated associations with the ACFT phenotypes and scores. As shown in **Figure 2** and **Figure 5**, regression models that employ these biological priors consistently outperform the control models that rely on equivalent regression algorithms (sPLSR) applied to the entire set of measured features for dimensionality reduction. This dimensionality reduction strategy proves particularly beneficial for analyzing datasets of modest effect size for discrimination of the relevant outcome by individual variables. PhenoMol allows for identification of groups of features (i.e. small molecules, proteins, transcripts) linked by common biology that reinforce one another to predict the outcome of interest in aggregate.

The network-based analysis in PhenoMol also provides a robust biological framework for data interpretation. Over-representation analysis (ORA) of the PN and PNM as summarized in **Figure 6** allows us to identify the most significant biological pathways associated with the ACFT outcome of interest. The approach to the biological annotation of the PN and PNMs presented here has several advantages over traditional GSEA approaches.^38,39^ First, the PNM integrates the omics measurements (i.e., metabolomics, proteomics and transcriptomics) through the interactome of known molecular relationships. Second, a binary outcome is not required to be defined to perform the ORA. Finally, the generation of the networks by the PCSF algorithm introduces Steiner nodes, that are linked in interaction space to the prize nodes but were not measured. These Steiner nodes help identify relevant biology to understand the PNM specific ensemble models.

The top biological annotations and the largest contributors to the PhenoMol models of ACFT are the complement and coagulation cascade (PNM 3) and primary bile acid biosynthesis (PNM 2). The complement system, which plays a role in both innate and adaptive immune responses, typically serves as the body’s first line of defense following infection. ^40^ With regard to the current work, recent studies suggest that the complement cascade is broadly upregulated following exercise^41^ and may play a role in regeneration and injury responses. ^42,43,44,45^ Notably, using a mouse muscle regeneration model, Zhang et al. identified a role for complement C3a in the recruitment of monocytes into injury sites and subsequent promotion of regeneration. ^46^

With respect to bile acids, which are involved in lipid absorption and liver energy metabolism, emerging studies in mice and humans support a role for bile acids in the regulation of skeletal muscle mass. ^47^ Other studies have demonstrated that bile acids serve as a signal governing muscle protein synthesis, energy metabolism, and regenerative capacity via the nuclear exporter FXR and the membrane transporter TGR5. ^47,48^ Additionally, we note that tauroursodeoxycholic acid, a bile acid metabolite identified in this study as being higher in the >540 ACFT cohort has also been shown in a mouse model to prevent dexamethasone induced muscle atrophy. ^49^

Arginine and proline metabolism (PNM 4) are also known to impact athletic performance through physiological and metabolic mechanisms. ^50^ Nitric Oxide Synthesis (NOS) enzymes catalyze the production of Nitric Oxide from L-Arginine. Nitric Oxide (NO) has been implicated in increased blood flow and improvement of muscle contraction, the kinetics of gas exchange, and even the reproduction of mitochondria. The signaling molecule is known to promote relaxation of vascular smooth muscle and ensuing dilation, which may favorably impact blood flow and augment mechanisms contributing to skeletal muscle performance.^51^ Additionally, there has been an athletic performance supplement industry built upon the promise of enhanced performance through the consumption of these metabolites and soluble precursors, yet the impact of these compounds is still under study.

Urea, which is a product of protein breakdown and degradation, also plays an important role in athletic performance as previously reported.^52,53^ The urea cycle, which is part of the Arginine and proline metabolism pathway (PNM 4), breaks down toxic ammonia generated from exercise-induced protein degradation and produces urea. Ammonia has been linked to physical fatigue.^52^ Further, the expression of the urea cycle enzymes are tissue-specific with higher levels found in skeletal muscle.^54^

Other noteworthy molecular differentiators of high ACFT performance included metabolites of the tryptophan-kynurenine pathway (PNM 1). Tryptophan is an essential amino acid and its breakdown via the kynurenine pathway accounts for much of its catabolism into several bioactive metabolites including kynurenine, kynurenate, quinolinate, and picolinate that have pleiotropic effects in several tissues. It has also been reported that in skeletal muscle, increased exercise elicits a PGC1α-mediated conversion of kynurenine into kynurenic acid, reducing the likelihood of depression and increasing resiliency under conditions of chronic stress.^55,56^

Recognizing the regulating role of bile acids, arginine, proline, urea, and tryptophan also offers the potential of using these natural products not only as a diagnostic tool to monitor or predict performance, but also as a therapeutic or prophylactic to enhance muscle performance.

In summary, PhenoMol was able not only to predict performance scores in candidates but also to identify biomolecular pathways and processes which may reveal the biological mechanisms associated with the differences between high- and low-performers. This approach opens the door not only to the design of new molecular assessment tools for human performance, but also avenues for tailored training, nutrition and/or supplements to achieve and maintain peak performance. The PhenoMol pipeline is a generalizable and efficient tool to make the best use of the rapidly expanding array of -omic, molecular and phenotypic assays from small cohorts, allowing studies of unique populations (e.g., elite athletic/military performance, rare diseases). The approach may also be leveraged in larger population studies where the depth and breadth of profiling employed here may be prohibitive, utilizing and refining the tools best suited to addressing the larger populations.

## Supporting information

multiple supplementary files

## SUPPLEMENTARY INFORMATION

The online link for supplementary info is provided here.

## ACKNOWLEDGEMENTS

This material is based upon work supported by the United States Air Force and Defense Advanced Research Projects Agency (DARPA) under Contract No. FA8650-19-C7945 (Distribution Statement “A”: Approved for Public Release, Distribution Unlimited). Any opinions, findings and conclusions or recommendations expressed in this material are those of the author(s) and do not necessarily reflect the views of the United States Air Force, the Department of Defense, or the U.S. Government. Our sincere gratitude to the US Military Academy cadets at West Point, who during the Spring of 2021 volunteered their times, provided valuable insights, and supported this study. Special thanks to COL Mark Reed, COL Phil Dacunto, and the West Point Department of Geography and Environmental Engineering for use of classroom space to conduct the data collection; COL Nicholas Gist, Todd Crowder, Jesse Germain, and the West Point Department of Physical Education for the use of their laboratory space and instrumentation and supporting the participation of the Sandhurst Black and Gold teams; The West Point Department of Military Instruction command team and Victor Castro, Ed McElvaney, and Gordon Cooke for the use of the West Point Simulation Center; The Office of the Dean of the Academic Board and the entire leadership team at the US Military Academy for their unconditional support and use of the facilities at the US Military Academy at West Point.

We are extremely thankful to Eric Van Gieson, Gopal Sarma, Adam Willis, Chris Kegelman and Lucas Veillon from DARPA; Christine Morton, Diane Minas, Peter Dafniotis, Natalie Guay and Martin Brown from GE HealthCare Technology & Innovation Center; Amber Hill, Erica Sampson, Arnyah Brown Countess, James Kubricht, Donald Hamilton and Habib Abi-Rached (from GE Research); Taisha Joseph, Moti Zwilling, Yu-Xin Xu, Stanislav Yuri Tsitkov and Matthew J. Leventhal (Massachusetts Institute of Technology) for their continuous advocacy, support, encouragement and valuable discussions. Special thanks to our commercial collaborators at Metabolon, Diagenode, AxisPharm, Eve Technologies, and Bruen Medical Partners for performing various molecular assays and phlebotomy services required in this study. Crystal Jaing, Jeff Drocco and their team at Lawrence Livermore National Lab are thanked for conducting independent quality assurance and quality control (QA/QC) analysis of all the phenotypic and molecular measurements presented in this paper. We are grateful to Adam Irvin and Sherrie Pryber from Air Force Research Labs; and Karen Peck from the US Military Academy at West Point for all their support and valuable insights to fulfill the IRB/HRPO requirements in support of the human subject study presented here.

## AUTHOR CONTRIBUTIONS

A.A. co-lead this work, conceived, designed and contributed to the execution of the human subject study at West Point, and drafted the manuscript. J.G. developed the bioinformatics framework, analyzed data, produced figures and contributed to manuscript writing. MM contributed to data analysis and manuscript writing. A.A.B. contributed to phenotypic assay design, figure production and manuscript writing. F.G. and N.H.H., contributed to discussions and manuscript writing. K. J. O., K. W., N.B. and G.F. contributed to the design and execution of the human subject study at West Point as well as discussions and manuscript writing. E. M., B.M.D., E.R.L., C.S., S.P., A.C., L.S., C.C., R.L., G.S.K., E.D.W. developed the SOPs for biospecimen sampling and processing, contributed to execution and collection of molecular measurements. P.T. J.W., O.B., R.G.S., T.H. collected the phenotypic data. S.S., J.J., S.C.E., R.X., S.C. and N.H. contributed to the analysis of the data and implementation of the bioinformatics pipeline. E.F. and L. M., co-lead this work and contributed to the development of the analytics framework and manuscript writing.

## METHODS

### KEY RESOURCES TABLE

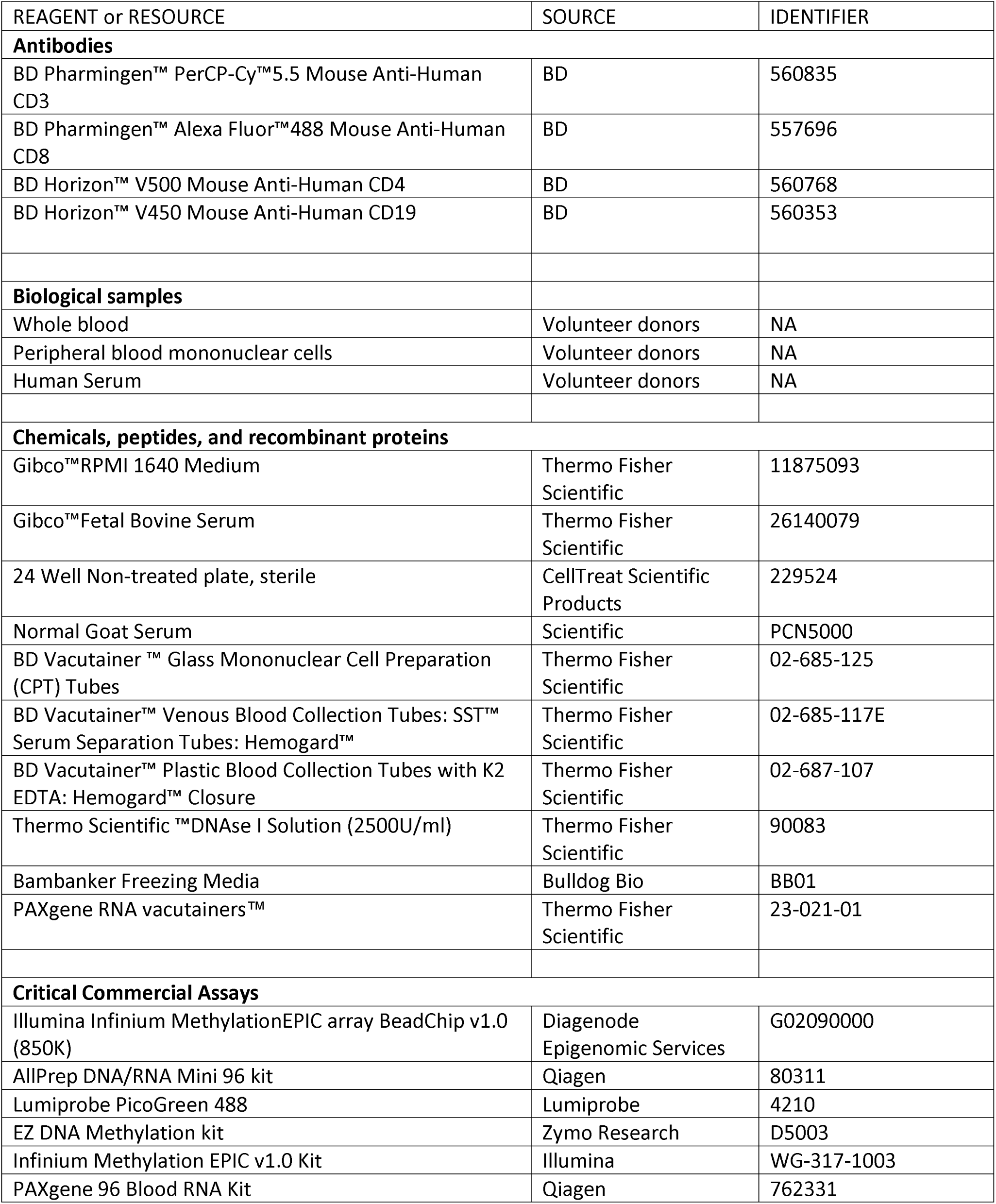
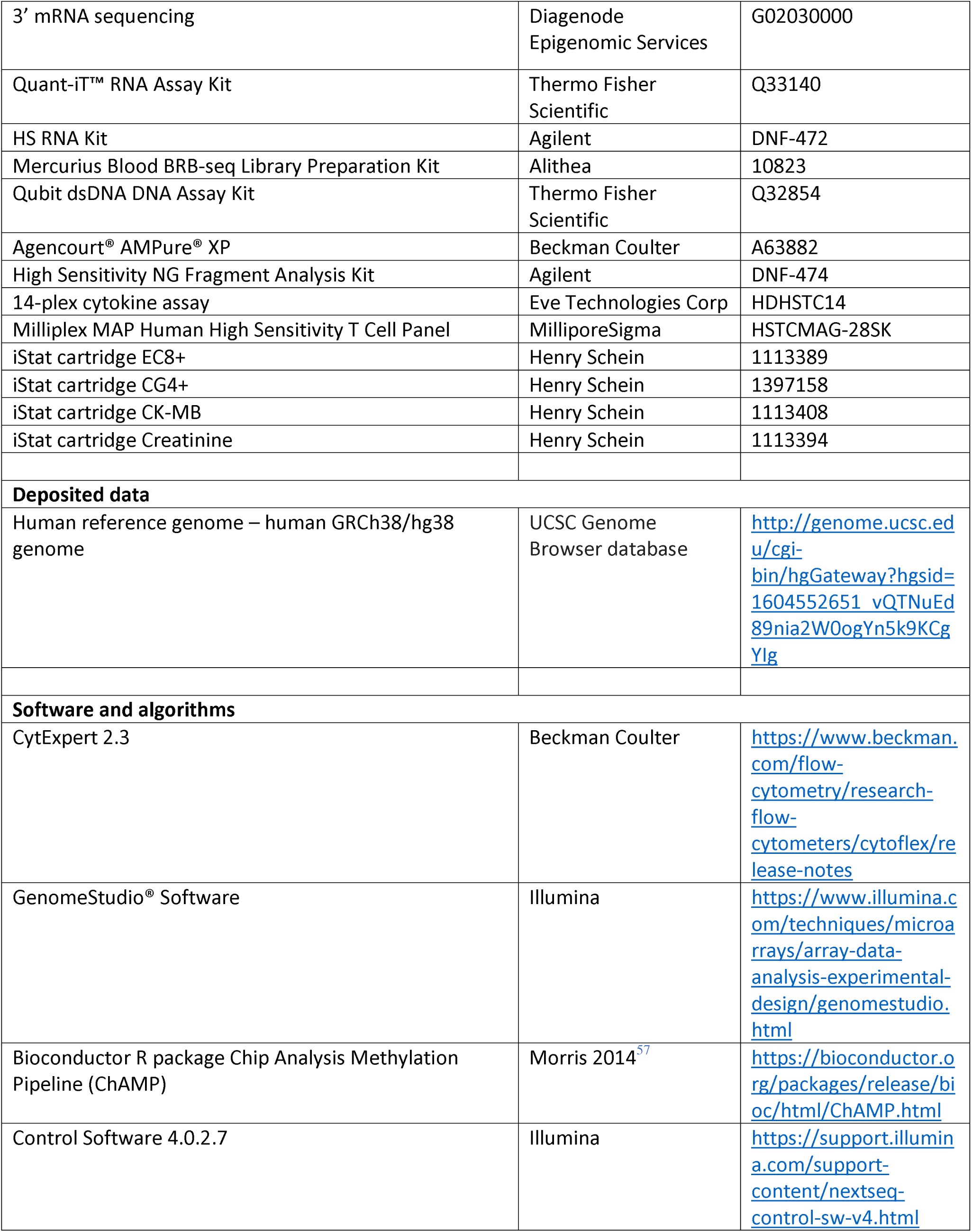
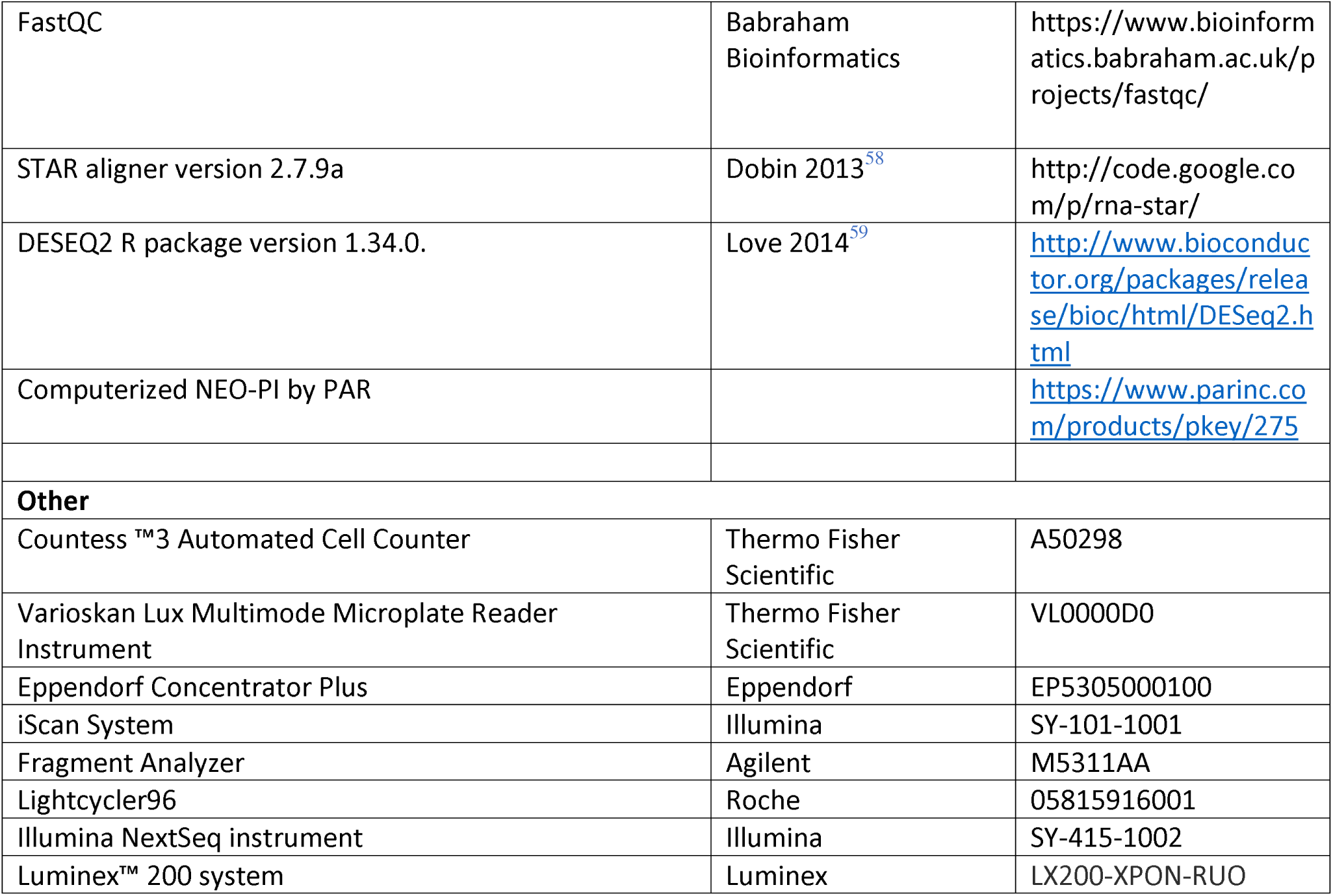

### RESOURCE AVAILABILITY

#### Lead Contact

Further information and requests for resources should be directed to and will be fulfilled by the Lead Contact, Azar Alizadeh.

#### Data and Code Availability

Codes will be placed on https://github.com/GE-Bio. Data may be available by request after review by the United States Military Academy Institutional Research Committee and is dependent on the type of request. Requests can be made at. The review will determine the risks to personnel for data sharing and is de-pendent on the type of request.

### EXPERIMENTAL MODEL AND SUBJECT DETAILS

#### Participant Recruitment and IRB Consent

71 healthy male (age 21±1 years) and 15 healthy female (age 20±1 years) cadets from the United States Military Academy (USMA) in West Point, NY were enrolled in this study. Participation in this research was voluntary and the 86 subjects provided written informed consent forms under the study research protocol FWR20200062H approved by the Air Force Research Laboratory (AFRL) Institutional Review Board / Human Research Protection Office (IRB/HRPO). All USMA cadets participating in military training and competition events at West Point were included in the recruiting and enrollment plan. Cadets on academic probation or limiting medical profiles, as well as cadets enrolled in other human performance research studies using wearable devices that conflicted with this study were excluded. Study participants consented to make their records from the Academy Management System (AMS) available to the research team in a coded format (i.e., with all personally identifiable information removed). The AMS records included information on cadet’s academic, military, and physical fitness assessments. Study participants provided demographic (such as gender, age, height, weight) and general health status information. Study participants consented to make their de-identified data open access for research purposes.

### METHODS DETAILS

#### Study Design

Immediately upon signing the consent form and enrolling in the study, participants received wearable devices, including a Garmin Fenix 6-series Watch/Heart rate monitor strap and an Oura ring (Generation 2), and were asked to wear them for the entire study duration of up to 3 months. These devices allowed for continuous monitoring of heart rate, respiration rate, and sleep patterns on a continuous basis. Each subject participated in up to five separate data collection sessions, each lasting approximately 3 hours.

#### Prescribed Exercise Protocol and Cardiopulmonary Performance Testing

Prescribed stationary cycling exercise trials completed through the course of the study were defined based on individual participant’s VO_2_max values. During the first trial session, each participant was asked to rate their familiarity with cycling as an exercise and overall fitness on a scale from 1-3. Based on their response, a power ramp rate was determined between 20 and 40 Watts/min. Each participant wore a Garmin Fenix 6 fitness tracking watch linked via ANT+ wireless to a Garmin HRM-Pro heart rate strap and Favero Asioma Uno power meter pedals mounted on a Monark 894E stationary cycle. After bike-fitting and warm-up (3 min. at 50W) each participant performed a ramped exercise profile (20, 30 or 40 W/min via weight addition) while monitored by a COSMED Fitmate Pro metabolic cart. The ramped exercise session was ended either by the participant at volitional exhaustion or by the study staff after peak VO_2_ had been reached. Based on the O_2_ consumption curve generated during this test, a prescribed power value was chosen at 70% of VO_2_max and used for all future exercise trials. During subsequent exercise trials, each participant performed 20-minute exercise trials with the same continuous monitoring of heart rate, power and cadence followed by blood draws or other assays. All participants were informed that exercise trials could be ended by either the participant or the study staff at any time based on discomfort, pain, or apparent dizziness.

#### Body Composition Testing

Participant’s body composition (body fat percentage, total and sectional lean muscle mass, etc.) and water distribution (intra/extracellular water) were measured on an InBody 770 analyzer through four electrode tissue impedance. Participants removed their socks and shoes to allow contact with foot electrodes and grasped hand electrodes to complete the circuit. An internal scale measured body mass, and impedance analysis was carried out automatically through the hand and foot electrodes to isolate the contributions of each limb and the torso to each variable.

#### Sleep and Daily Activity Tracking

Study participants were provided (at their discretion) with off-the-shelf fitness and sleep monitoring devices including an Oura ring (Generation 2) and a Garmin Fenix 6 or 6S watch (loaded with specific workout profiles for study activities). The Oura rings were worn only at night to monitor sleep, while the Garmin watches were worn continuously at the discretion of the participant. The Oura ring measures heart rate, body temperature and activity and outputs a variety of sleep metrics including time per sleep phase, heart rate variability, and resting heart rate. The Garmin watch directly measures heart rate and movement, has wireless connectivity with the power meter pedals for exercise trials, and algorithmic outputs including sleep time (by phase), daily movement (steps, flights of stairs, etc.) and rest time among others. As these devices were provided and used at the discretion of the study population, some participants had limited or no recorded data, while others only used one device or the other. In cases where sleep metrics were available from both devices, the Oura ring was used. In cases where only Garmin data was available, comparable metrics were used, and for participants lacking both, imputed data was used.

#### Personality and Behavioral Analysis by NEO PI-3

The Revised Neuroticism-Extraversion-Openness Personality Inventory (NEO PI-R) is a personality inventory that examines a person’s big five personality traits (openness to experience, conscientiousness, extraversion, agreeableness, and neuroticism). In addition, the NEO PI-R reports on six subcategories of each of these personality traits, called facets. The development of the Revised NEO PI-R began in 1978 with the publication of a personality inventory by Costa and McRae.^60^ The 2005 updated version of this personality inventory, referred to as NEO PI-3, contains updated vocabulary that could be understood by adults of any education level. The computerized NEO PI-3 version by PAR, consisting of 240 items with detailed facet scores, was administered in this study. The test was taken by candidates on laptops/tablets without any time limitations (average time to complete the test was approximately 35 minutes). The results were scores, and reports generated by PAR. In this study the t-scores for the main personality traits and the sub-category facets were used.

#### Cognitive Performance Analysis by Raven Progressive Matrices and Wisconsin Card Sorting Test

*Raven’s Progressive Matrices (RPM)*: The RPM is a non-verbal test for fluid intelligence, including problem solving and adaptive reasoning. The RPM test comprises 60 multiple choice questions, listed in order of increasing difficulty. This test was administered on a tablet.

##### Wisconsin Card Sorting Test (WCST)

The WCST is a neuropsychological test to measure cognitive flexibility or set shifting. The test measures the ability to display flexibility and adaptation in the face of sudden change in reinforcement and probes multiple cognitive functions such as working memory, attention, executive control, and visuo-spatial processing, which have been demonstrated to be affected by external stressors. The test was introduced by David A. Grant and Esta A. Berg in 1948. Four cards are on the table and one stimulus card is presented to the participant. The participant is told to match the cards, but not how to match (color, symbol, or number). When they make a guess to match the cards, they are told if they are right or wrong. The participant will eventually guess the matching criterion and use this criterion in future trials. Eventually, without notice to the participant, the matching criterion is changed, and the participant needs to react and determine the new matching criterion. The test will be carried out on a computer or tablet in each trial and the results will be compiled by the software including number, percentages, and percentiles of trials, errors, and perseverative errors. The test provides insight in frontal lobe function such as strategic planning, organized searching, utilizing feedback to adapt to shifting conditions, directing behavior toward achieving a goal, and modulating impulsive response. We ran a computerized version of the tests and conducted 2 trials at rest to baseline the response and measure learning effects of each subject. The study participants were asked to pedal for 20 min at 70% of their VO2 max on a stationary bike. Immediately after exercise, the cadets transitioned to one last WCST trial to measure the changes in their cognitive performance post physical stress.

#### Blood Collection and Sample Preparation

Venous blood was collected via venipuncture in the antecubital fossa (inner elbow) from participants at baseline (before exercise) as well as approximately 2-5 min, 10-20 min, and 25-40 min post-exercise. Specimens were immediately processed after collection. For separation of serum, blood was collected in a gold top 5 mL SST vacutainer™ (Cat# 367989, Becton Dickinson), inverted 5x to mix, and allowed to clot at room temperature for 15 min. Vacutainers™ were then spun at 1500xg for 15 min at room temperature. The upper serum layer was pipetted off, aliquoted and immediately frozen on dry ice. For plasma separation, blood was collected in a purple top 4 mL K_2_EDTA vacutainer™ (Cat# 368047, Becton Dickinson) and inverted 8x to mix. At the baseline blood draw, an aliquot of whole blood was taken from the K_2_EDTA vacutainers™ and stored in 1.5 mL Eppendorf tubes at 4°C for 4 to 16 hours prior to hematology analysis. For plasma collection, the K_2_EDTA vacutainers™ were spun at 1500xg for 15 min at room temperature. The top plasma layer was pipetted off, aliquoted and immediately frozen on dry ice. Blood for iSTAT analysis was collected in a green-top 2 mL Lithium Heparin vacutainer™ (Cat# 366664, Becton Dickinson) and inverted 4x to mix. Blood was collected in 8mL CPT vacutainers™ (Cat# 362753, Becton Dickinson) for separation of peripheral blood mononuclear cells (PBMCs) from whole blood. A 15 min centrifugation was performed (1500xg) to separate the mononuclear cells from the erythrocytes and granulocytes. The PBMCs were washed in PBS (5 min, 250xg) at room temperature to deplete residual contaminants such as platelets and erythrocytes and to dilute the plasma. PBMCs were then resuspended in 1mL PBS and counted on a Countess™3 automated cell counter. Aliquots of PBMCs were flash-frozen as dry pellets for DNA methylation analysis. For immunophenotyping, PBMCs were resuspended in 1mL Bambanker cryopreservation reagent (Cat# BB01, Bulldog Bio) and immediately stored in a Mr. Frosty freezing container on dry ice then transferred to liquid nitrogen. Whole blood was stabilized in 2.5mL PAXgene RNA vacutainers™ (Cat# 762165, Becton Dickinson) for 4-16 hours at room temperature then stored at -20°C prior to RNA transcriptome analysis.

#### DNA Methylation Analysis from Peripheral Blood Mononuclear Cells

##### DNA extraction and bisulfite conversion

The Illumina Infinium MethylationEPIC array BeadChip v1.0 (850K) was carried out by Epigenomic Services at Diagenode (Cat# G02090000) using bulk PBMC dry pellets. In brief, the PBMCs were thawed on ice and extracted in a randomized order using the AllPrep DNA/RNA Mini 96 kit (Cat# 80311, Qiagen) across four plates. The plates were extracted according to an adapted protocol from the manufacturer to increase DNA recovery. The approximate elution volume of the extracted gDNA was 70 µL. After extraction, the samples’ gDNA were quantified using the Lumiprobe PicoGreen 488 (Cat# 4210, Lumiprobe) on a Varioskan Lux Multimode Microplate Reader Instrument (Cat# VL0000D0, Thermo Fisher Scientific). Samples that had a concentration above 10 ng/µL went into the bisulfite conversion step with a gDNA input of 500 ng. Samples that had concentrations below 10 ng/µL went through a concentration step using the Eppendorf Concentrator Plus (Cat# EP5305000100, Eppendorf) and entered the bisulfite conversion with a gDNA input of up to 500 ng. During the bisulfite conversion step, the gDNA was deaminated utilizing the EZ DNA Methylation kit (Cat# D5003, Zymo Research) in accordance with Illumina’s preferred deamination protocol.

##### DNA sequencing, quality control, and data processing

The samples were loaded on the Infinium MethylationEPIC v1.0 Kit (Cat# WG-317-1003, Illumina) and inserted into an iScan System (Cat# SY-101-1001, Illumina) for an array scan as per manufacturer protocol. Samples that entered the bisulfite conversion workflow with gDNA inputs below 250 ng were excluded from downstream analysis. The GenomeStudio® Software was next used to examine the built-in Infinium controls and understand the data quality. The Illumina Methylation Module user guide has a Controls Dashboard section that exhaustively explains the analysis process and can be reviewed for additional details. In brief, controls were included for staining, extension, hybridization, target removal, bisulfite conversion efficiency, and non-specific primer extension, as well as negative controls lacking CpG dinucleotides to define the background signal intensity. Non-polymorphic controls were used to assess the sample quality and overall assay performance. After completing the Array QC steps using the built-in controls, the dataset was imported into the Bioconductor R package Chip Analysis Methylation Pipeline (ChAMP) for further QC and bioinformatic analysis. The additional QC steps evaluate each samples’ proportion of failed probes, distance clustering between samples (using all probes), methylation distribution plots, and multidimensional scaling. The primary steps in the pipeline include filtering, normalization with BMIQ, singular value decomposition analysis, and batch effect correction with the COMBAT algorithm. After the filtering and processing, the final dataset consisted of 741,605 probes with methylation values. The final data matrix was exported into a csv containing the methylation values for all samples for the 741,605 probes and was integrated into the omics integrator pipeline.

#### RNA Sequencing from Stabilized Whole Blood

Transcriptome analysis, including RNA extraction and 3’ mRNA sequencing, was conducted by RNA Seq Services at Diagenode (Cat# G02030000).

##### RNA Extraction

The whole blood PAXgene vacutainers™ were thawed at room temperature and underwent RNA extraction using the PAXgene 96 Blood RNA Kit (Cat# 762331, Qiagen) in a randomized order. The extracted total RNA was eluted in 80 uL and quantified using the Quant-iT™ RNA Assay Kit (Cat# Q33140, Thermo Fisher Scientific) on a Varioskan (Cat# VL0000D0, Thermo Scientific). The sample quality and integrity were evaluated by calculating the RNA Quality Number (RQN) using HS RNA Kit (Cat# DNF-472, Agilent) on the Fragment Analyzer (Cat # M5311AA, Agilent).

##### Library preparation for 3’ mRNA sequencing

The total RNA input for library preparation was 50 ng for each sample. The Mercurius Blood BRB-seq Library Preparation Kit (Cat# 10823, Alithea) was next used to perform polyA selection, cDNA synthesis, and adapter ligation using the manufacturer’s protocol. Globin depletion is integrated to the kit’s library preparation technology, whereby the cDNA synthesis occurs in the presence of oligos that block the reverse transcription of globin mRNA. The cDNA yield of each pool was quantified using the Qubit dsDNA HS DNA Assay Kit (Cat# Q32854, Thermo Fisher Scientific) and approximately 8 ng of pooled cDNA was used for library preparation. To determine the number of amplification cycles required to achieve an appropriate library size, a small volume of the purified adaptor-ligated cDNA was evaluated by qPCR on a Lightcycler96 (Cat# 05815916001, Roche) as per manufacturer’s protocol. Adapter sequences were used for the amplification to pool two microtiter plates per lane for sequencing in a total of 3 lanes. After amplification and purification using Agencourt® AMPure® XP (Cat# A63882, Beckman Coulter), the libraries were again quantified using the Qubit™ dsDNA HS DNA Assay Kit). Next, a High Sensitivity NG Fragment Analysis Kit (Cat# DNF-474, Agilent) was used to evaluate the mean cDNA size and size range on a Fragment Analyzer™ (Cat # M5311AA, Advanced Analytical Technologies, Inc).

##### 3’ mRNA sequencing and data processing

The pooled libraries – split into 5 sequencing pools – were sequenced on an Illumina NextSeq instrument (Cat# SY-415-1002, Illumina) that used Control Software 4.0.2.7 in paired-end mode. Read 1 sequence (forward read) contains the barcode sequence and UMIs. Read 2 sequence (reverse read) contains index sequence and the cDNA sequence. FastQC was used to control the quality of the raw sequenced reads. ^61^ Demultiplexing was performed using Bcl2fastq software, then STAR aligner version 2.7.9a was used to align the reads to the human GRCh38/hg38 reference genome (UCSC). The primary sequencing and alignment statistics evaluated per pool include total read pairs, uniquely aligned read pairs and ratio, multimapping read pairs and ratio, and unmapped read pairs and ratio. The number of detected genes was plotted on a per sample basis, with coverage, defined as unnormalized number of counts, examined at more than 0 (detected), more than 10 and more than 100. The presence of outliers was evaluated using DESeq2 from the DESEQ2 R package version 1.34.0. ^59^ DESeq2 was also used create a data matrix of normalized counts for all genes. Four data matrices were created with the permutations of unnormalized and normalized counts against ensemble gene ID or gene name for all samples and exported into csv samples. These data matrices were then integrated into the omics integrator platform for additional analysis.

#### Untargeted Proteomics Analysis from Plasma

Untargeted proteome analysis was conducted at AxisPharm LLC

##### Sample Preparation

Previously aliquoted K_2_EDTA plasma samples were thawed on ice and randomized for subsequent analysis. After thawing, 2 µL samples were mixed with 100µL 6M guanidine containing a standard protein cocktail boiled for 5 min then chilled to -20°C for 10 min. Samples were then extracted with methanol (1mL) and incubated at -80°C for 1 hour, followed by centrifugation. After liquid removal, samples were resuspended in 200 µL 8M urea/0.2M NH5CO3 and incubated at -20°C overnight. 50 µL aliquots of each sample were mixed with 50 µL of 50mM NH_5_CO_3_ and 0.2 µg end proteinase LysC then incubated at 37°C for 2-3hrs. 100 µL of 50mM NH_5_CO_3_ with 0.1µg Trypsin was added to each sample and incubated at 37°C overnight, followed by 1 hour at 42°C. 1mL of 0.1% formic acid was added to each sample, vortexed and the resulting solution was added to a Solid phase extraction plate (8E-S100-AGB Strata™-X 33 µm Polymeric Reversed Phase, 10 mg / well, 96-Well Plates). Each well’s contents were activated with 100% acetonitrile, equilibrated, and washed with 0.2% formic acid, eluted with 40% acetonitrile, dried and resuspended in 200µL 0.5% formic acid, 5% acetonitrile, 94.5% H2O. The Trypsin-digested peptides were analyzed by ultra-high pressure liquid chromatography (UPLC, Thermo Dionex UltiMate™ 3000 RSLC nano System) coupled with LC-MS/MS tandem mass spectroscopy (Orbitrap Fusion Lumos, Thermo Fisher Scientific) using nano-spray ionization. The instrumentation used was an interfaced with nanoscale reversed phase UPLC

##### Sample Analysis

Samples were loaded onto a precolumn (volume less than 15 µL, Cat# 164564-CMD, Thermo ACCLAIM PepMap 100) at a 7 µL/min flow rate for 5 minutes followed by the analytical run using a 25 cm, 75-micron ID glass capillary packed with 1.7 µm C18 (130) BEHTM beads (Waters Corporation). Peptides were eluted from the C18 column into the mass spectrometer using a linear gradient (5–100%) of acetonitrile (ACN) at a flow rate of 395 μL/min for 2 hours. The buffers used to create the ACN gradient were Buffer A (98% H2O, 2% acetonitrile, 0.1% formic acid) and Buffer B (20% H2O, 80% acetonitrile, 0.1% formic acid).

Mass spectrometry analyses were carried out in two steps. First, a MS1 survey scan was made with the using the Orbitrap detector (mass range m/z: 400-1500 using quadrupole isolation, 60000 resolution setting, spray voltage of 2400 V, ion transfer tube temperature of 290°C, AGC target of 400000, and maximum injection time of 50 ms). The survey scan was followed by data-dependent scans (top speed for most intense ions, with charge state set to only include +2-5 ions and 20 second exclusion time), with ion selection (minimal intensities of 50000 or greater), and HCD Collision Energy of 30%. The fragment masses were analyzed in the ion trap mass analyzer (with the ion trap scan rate set to turbo, first mass m/z = 100, AGC Target of 5000 and maximum injection time of 35ms). Protein identification was carried out using the Peaks Studio 8.5 software package (Bioinformatics Solutions Inc.)

#### Untargeted Metabolomics Analysis from Plasma

Untargeted metabolome analysis was conducted using the HD4 Platform at Metabolon Inc.

##### Sample preparation

The frozen K_2_EDTA plasma samples were thawed on ice then extracted with methanol under vigorous shaking for 2 min (Glen Mills GenoGrinder 2000) to precipitate protein and dissociate small molecules bound to protein or trapped in the precipitated protein matrix, followed by centrifugation to recover chemically diverse metabolites. The resulting extract was split into five fractions: two fractions were used for positive ion electrospray (ESI) reverse phase (RP)/UPLC-MS/MS analysis using two separate methods, fraction 3 was used for negative ion mode ESI for analysis by RP/UPLC-MS/MS, fraction 4 was used for negative ion mode ESI analysis by HILIC/UPLC-MS/MS, and the fifth fraction was saved as a backup sample if repeat analysis was needed. Samples were placed briefly on a TurboVap® (Zymark) to remove organic solvents. The sample extracts were stored overnight under nitrogen before preparation for analysis.

##### Quality Control (QC)

Quality control samples were analyzed in concert with each set of experimental samples. These include: 1) technical replicate samples derived from a pool of well-characterized human plasma (MTRX) or, alternatively client matrix generated by combining a small portion of each (non-plasma) experimental sample (CMTRX); 2) extracted water samples (process blanks) and solvent blanks; and 3) a cocktail of QC standards, carefully chosen to avoid interference with the measurement of endogenous compounds. The QC samples enabled instrument performance monitoring and aided with chromatographic alignment. Instrument variability was determined by calculating the median relative standard deviation (RSD) for the standards. Overall process variability was determined by calculating the median RSD for all endogenous sample-derived metabolites (i.e., non-instrument standards) present in each of the pooled MTRX or CMTRX technical replicate samples. Experimental and QC samples were randomly interspersed.

##### Ultrahigh Performance Liquid Chromatography-Tandem Mass Spectroscopy (UPLC-MS/MS)

Measurements were acquired on a Waters ACQUITY ultra-performance liquid chromatography (UPLC) and a Thermo Scientific Q-Exactive high resolution/accurate mass spectrometer interfaced with a heated electrospray ionization (HESI-II) source and an Orbitrap mass analyzer operated at (35,000 mass resolution). The dried sample extracts were reconstituted with standards in solvents compatible with each method. Each reconstitution solvent contains a series of standards at fixed concentrations to ensure injection and chromatographic consistency. Four methods were employed to broadly detect all metabolite compounds. Hydrophilic compounds were gradient-eluted from a C18 column (Waters UPLC BEH C18-2.1×100 mm, 1.7 µm) gradient using water and methanol, containing 0.05% perfluoropentanoic acid (PFPA) and 0.1% formic acid (FA) then analyzed using positive ion electrospray (ESI) reverse phase (RP)/UPLC-MS/MS. Hydrophobic compounds were eluted with using methanol, acetonitrile, water, 0.05% PFPA and 0.01% FA then analyzed using positive ion electrospray (ESI) reverse phase (RP)/UPLC-MS/MS at an overall higher organic content. The third aliquot was analyzed using negative ion mode ESI for analysis by RP/UPLC-MS/MS using a separate dedicated C18 column, gradient-eluted from the column using methanol and water, with 6.5mM Ammonium Bicarbonate at pH 8. The fourth aliquot for polar compounds analyzed via negative ion mode ESI analysis by HILIC/UPLC-MS/MS negative ionization following elution from a HILIC column (Waters UPLC BEH Amide 2.1×150 mm, 1.7 µm) using a water/acetonitrile gradient with 10mM Ammonium Formate, pH 10.8. The MS analysis system alternated between survey scanning and MS and data-dependent MSn scans using dynamic exclusion. The scan range varies varied slightly between methods covers approximately 70-1000 m/z.

##### Data Extraction and Compound Identification

Raw data underwent extraction, peak-identification and QC processing using Metabolon’s in-house hardware and software. These systems are built on a web-service platform utilizing Microsoft’s .NET technologies, which run on high-performance application servers and fiber-channel storage arrays in clusters to provide active failover and load-balancing. Compounds were identified using Metabolon’s MS comparison to a library of more than >4500 commercially available pure standards and recurrent structurally unnamed biochemical species. Metabolon’s library is based on authenticated standards and contains the retention time/index (RI), mass mass-to to-charge ratio (m/z) and underlying chromatographic data (including MS/MS spectral data) on all molecules present in the library. All biochemical identifications are based on three criteria: retention index within a narrow RI window of the proposed species, accurate mass match (+/- 10 ppm), and MS/MS forward and reverse scores. MS/MS scores are based on a comparison of the ions present in the experimental spectrum to ions present in the library entry spectrum. The use of all three measures enables high confidence in the molecular species assignments.

#### Hematology, Metabolic Panel and Blood Gas Analysis from Whole Blood

Whole blood samples stored at 4°C were gently flicked to mix immediately prior to analysis on the Sysmex XN-330, providing a complete blood count with diff. The parameters measured include WBC, RBC, HGB, HCT, MCV, MCH, MCHC, PLT, RDW-SD, RDW-CV, PDW, MPV, P-LCR, PCT, NEUT#, LYMPH#, MONO#, EO#, BASO#, NEUT%, LYMPH%, MONO%, EO%, BASO%, IG#, and IG%. Data outside the normal range were flagged and recorded.

Whole blood collected in Lithium Heparin vacutainers™ was added to test-specific i-Stat cartridges (Henry Schein) 5-10 minutes after blood draw. 95 uL of blood was dispensed into a CG4+ cartridge to analyze lactate concentrations, 65 uL of blood was dispensed into EC8+ cartridge to analyze glucose, BUN, hematocrit and hemoglobin, 65 uL of blood was dispensed into a Creatinine cartridge, and 17 uL of blood was dispensed into a CK-MB cartridge to analyze Creatine Kinase MB concentrations.

#### Immunophenotyping from Cryopreserved Peripheral Blood Mononuclear Cells (PBMC)

Cryopreserved PBMC were thawed in a 37C water bath for less than 1 minute and resuspended in RPMI complete medium (10% FBS). ^62^ The cryopreserving solution was removed by spinning the cells at 200xg for 5 minutes. Cells were resuspended in 1 mL of complete RPMI and allowed to equilibrate at 37°C, 5% CO2, 95% humidity in a 24-well tissue culture non-treated plates for 1h. 50U/mL DNase was added to prevent cell aggregation. Cells were then counted and assessed for viability using a Countess™3 automated cell counter and up to 2×10^6^ viable cells were used per sample. Cells were blocked for 15 min with 10% normal goat serum, then were surface stained with a cocktail of fluorochrome-conjugated antibodies for CD3, CD4, CD8, and CD19 at predetermined optimal dilutions for 15 min at room temperature (see Table for antibody clones and conjugates). Cells were then washed with PBS and analyzed immediately on a flow cytometer (Beckman Coulter CytoFlex S). Data analysis was performed using CytExpert software version 2.3 for the percentage of cells expressing each surface marker within the live, singlet lymphocyte subpopulation.

#### Targeted Cytokine Analysis from Serum

Serum samples were thawed on ice, and subsequently plated and analyzed in a randomized order by Eve Technologies Corp (Calgary, Alberta) using their HDHSTC14 assay. The assay was a 14-plex kit utilizing the Luminex xMAP technology for the following analytes - GM-CSF, IFNy, IL-1β, IL-2, IL-4, IL-5, IL-6, IL-8, IL-10, IL-12p70, IL-13, IL-17A, IL-23, and TNF-α. The base kit is the Milliplex MAP Human High Sensitivity T Cell Panel (Cat# HSTCMAG-28SK, MilliporeSigma), and the samples were processed according to the manufacturer’s protocol. A Luminex™ 200 system (Luminex) was used to perform the multiplexing analysis. The data output included the fluorescence intensities raw data signal and observed concentration for the kits’ standards and the samples.

### QUANTIFICATION AND STATISTICAL ANALYSIS

#### Standard statistical analysis and data processing

R (version 4.2) was used to perform standard data processing, statistical testing, correlative analysis, plotting and figure generation. The non-parametric Wilcoxon rank sum test was used for all comparisons of two independent groups. Spearman’s rank correlation coefficient was computed to characterize the monotonic relationship between all two variable samples such as between a phenotype measurement and a molecular feature measurement.

#### PhenoMol computational tool

The PhenoMol models and results presented in this manuscript were generated using the PhenoMol version 1.0 computational tool running it from Jupyter Notebooks. PhenoMol version 1.0, can be downloaded from github (https://github.com/GE-Bio/PhenoMol) and currently requires python (https://www.python.org) version 3.10 and R (https://www.r-project.org) version 4.2 to be installed. The PhenoMol python files are dependent and utilizes methods from the following python packages:

- numpy version 1.26.4
- pandas version 2.1.3
- networkx version 3.2.1
- seaborn version 0.13.2
- SciPy version 1.11.4
- scikit-learn version 1.3.2
- pcst-fast version 1.0.10
- python-louvain version 0.16

The PhenoMol python files include calls to PhenoMol R-script files. These R script files require the mixOmics (version 6.22) R package available from Bioconductor. The PhenoMol computational tool runs a set of predefined steps for a standard PhenoMol analysis run. We will first summarize the steps followed by more detailed explanations on the key analytics.

1) **Setup and prepare data for analysis:** Define analysis run configuration parameters and create analysis run and results directories. Load raw phenotypic and omics data from comma-separated value (.csv) files. Process omics data that started by computing the median value measurement for each molecular feature across an individual’s blood draws and weekly events. Next features underwent a course filtering step and removed those features if they had missing values in more than 40% of the individuals or had no variation across the individuals.
2) **Perform k-fold stratification**: Define and create 5-folds of the data using the sklearn method StratifiedKFold from the python package scikit-learn. The StratifiedKFold method stratified subjects for high and low ACFT Total Points as defined by being above or below the median value of 553 for the cohort male population. The StratifiedKFold method uses an integer to initialize the pseudorandom number generator. We arbitrary selected the six integers, 123, 321, 456, 654, 789, 987 for six 5-fold cross validation analysis runs to effectively generate thirty unique training and test folds.
3) **Impute fold’s training data:** Within each fold, the training data (80% of the data) underwent imputation to deal with missing data. We selected the sklearn method KNNImputer to perform imputation for missing values using k-Nearest Neighbors where k was set to equal five neighbors. We performed this imputation within class of data type. Phenotypes, metabolites, proteins, transcripts, and DNA methylation were all treated as different classes for the purposes of nearest neighbors’ imputation.
4) **Generate prizes**: Raw prizes were generated using the imputed fold training data. We defined prizes as the absolute spearman correlation between a targeted phenotype or outcome and a molecular feature measurement. A p-value for the prize (i.e. spearman correlation) was computed and used to filter the prizes. We set the threshold p-value to 0.2 keeping prizes below this threshold value. We generated sets of prizes for the ACFT Total Points outcome and for the six ACFT events (i.e. targeted phenotypes) that included maximum deadlift, standing power throw, hand release pushups, sprint drag carry, leg tuck, and 2-mile run.
5) **Generate robust subnetworks:** Within the fold, a graph instance was created by loading in the interactome of prior data that consists of molecular feature nodes and edges between the nodes. An edge represents an interaction between the two nodes it connects with an associated cost of one minus the fractional confidence level in the interaction between the two nodes. Next, we generated independent robust subnetworks for the ACFT Total Points outcome and for the six ACFT event phenotypes using a modified Omics Integrator python script that was originally developed at the Fraenkel Lab at MIT. The original Omics Integrator python script file “grapy.py” was renamed “oi3_graph.py” within the PhenoMol tool. We modified the python script to utilize networkx version 3.2 and the Prize Collecting Steiner Forest (PCSF) algorithm implemented in the pcst-fast (version 1.0.10) python package. There are a set of hyperparameters that need to be set depending upon the context of the problem. Our objective is data dimensionality reduction via graph optimization. To this end we set the hyperparameter w, representing the cost of edges connecting the root dummy node to terminals, to unity. We set the hub penalty parameter g, which introduces a penalty to edges based on the degrees of the nodes they connect to zero. For the hyperparameter b, which scales prize values of the nodes relative to the edge costs, we utilized a tuning method that searched the b parameter space to generate a subnetwork of desired targeted size with the objective of balancing data reduction (number of terminal nodes) with network richness and interpretation (seek approximately equal number of Steiner nodes to terminal nodes). Tuning was performed using the Brent’s method implemented as optimize.brentq within the python SciPy package. We selected 128 terminal nodes to tune the hyperparameter b because it was smaller than the number of available measured metabolite or protein features thus promoting data reduction while at the same time generating networks that included a significant number of Steiner nodes. The Steiner nodes cannot be used to generate a model because they have not been measured, however they help with biological annotation and interpretation for the derived models. Furthermore, they can provide potential insights and hypothesis for future studies and features to target and measure. For the measured omics we have two broad categories of molecular features that included functional metabolites and proteins and genomic DNA methylation and RNA transcripts. We desired models that incorporated both functional and genomic features in them. To achieve this balance, we introduced a fourth hyperparameter, r, that is the ratio of the tuned hyperparameter b when generating a subnetwork using only prizes for the genomic features (DNA methylation and RNA transcripts) divided by the tuned hyperparameter b when generating a subnetwork using only prizes for the functional features (metabolites and proteins). With the four defined or tuned hyperparameters (w, b, g, r), the next step was to generate a robust subnetwork using all the available functional and genomic prize data. This was accomplished by running the PCSF algorithm to generate a graph after first applying random gaussian noise to the edge costs of the interactome (via the numpy.random.normal method with the scale parameter set to 0.1). This was repeated one hundred times to generate a set of graphs. The set of nodes from the set graphs were then individually scored by the frequency of the individual node appearing in the one hundred generated graphs. The robust subnetwork is defined as the nodes that appeared in at least 20% of the generated graphs along with the edges from the interactome that interconnect these nodes. This entire process was repeated for the ACFT Total Points outcome and for the six ACFT event phenotypes.
6) **Generate principal network and principal network modules:** Within each fold, the principal network was generated by taking the union of the nodes from the robust subnetworks for ACFT Total Points outcome and the six ACFT event phenotypes. The edges for the principal network are defined by the interactome edges that interconnect the principal network nodes. Next Louvain clustering (python-louvain version 0.16) was used to subdivide the principal network up into principal network modules. Louvain clustering was performed one hundred times using a different random state start, and for each the number of clusters and the heterogeneity index of the clustered nodes is computed. The heterogeneity index is the Shannon Diversity Index divided by the log of the number of distinct clusters. The Shannon Diversity Index is the summation of the quantity of each cluster’s proportion multiplied by the natural log of the cluster’s proportion. The clustering instance with the fewest number of clusters is selected. If there are multiple choices, the one with the minimum heterogeneity index is selected reflecting a more equal proportion of nodes among each cluster.
7) **Build fold’s base expression axis models:** The Bioconductor R package, mixOmics (version 6.22) was used to perform Sparse Partial Least Squares Regression (sPLS-R) analysis. ^63^ Base PNM EA models were generated for the outcome, targeted phenotypes, and each of the fold’s PNMs by sPLS-R using the folds training data. A base model was generated for each individual PNM and the entire PN (labeled PNM0). Only features that were members of the PNM or PN were used as possible candidate features for sPLS-R. Furthermore, we applied the option to generate a base model using all available fold omics features independent of the PN and PNMs (labeled PNM-99) thus applying the sPLS-R directly to the data (i.e. control models). The PhenoMol tool utilizes the three methods tune.spls, spls, and perf from the mixOmics R package to perform sPLS-R. There are multiple configurable hyperparameters although we set them to the following. For tune.spls we used mean square error (MSE) for the accuracy measure. We used Mfold for the internal tune.spls validation with three folds and fifty repeats. For defining the maximum number of features to keep, we limited it to be less or equal to the number of fold training subjects divided by three. Following performance testing of spls models (perf method of mixOmics package), we applied a feature stability filter and only kept features that achieved a minimum stability of 80% on the resulting sPLS models.
8) **Generate fold’s ensemble models:** Ensembles of the PNM EA models were generated for the outcome and targeted phenotypes. The ensemble models are built from multiple individual base PNM EA models (i.e. base learners) and combined to make a single prediction from the average of the individual predictions giving equal weight to each base PNM EA model prediction. We set the option of the maximum number of individual base models in an ensemble model to be two. Ensemble models were generated for every unique combination of two base models. Furthermore, the code allows an individual PNM base model to be combined with the PN base model.
9) **Impute fold’s test data:** Within each fold, the test data (20% of the data) underwent imputation to deal with missing data using the same method for imputation of fold’s training data. For imputing the test data, we included the fold’s training subject data when submitting to the sklearn method KNNImputer to perform imputation for missing values using k-Nearest Neighbors.
10) **Generate fold’s model test statistics:** The fold’s base and ensemble models were applied using the inputted test data to generate predictions which were then evaluated against the test data’s actual measurements to compute the fold’s test statistics of MSE and R2.

#### Pathway and Gene Set Enrichment Analysis

The Bioconductor R package, hypeR (version 1.14) was used to perform over-representation analysis, also sometimes referred to as geneset enrichment analysis. The over-representation analysis was performed on the entire principal network and each of the principal network modules. Both terminal nodes (measured) and Steiner nodes (not measured) were provided as input to the over-representation analysis as well as the pathways (i.e. genesets). We performed the over-representation analysis on the Kegg Legacy pathways that we downloaded from the Molecular Signatures database.^64^ These downloaded pathways identified which gene IDs were members of which pathway. We then linked DNA methylation, RNA transcripts, and proteins to these gene IDs based on the Ensembl version 110 (www.ensembl.org). For metabolites, we used the metabolite-pathway assignments stored in the Human Metabolome Database (HMDB) version 5^65^ to link each metabolite with a HMDB ID to the Kegg Legacy pathways.

## REFERENCES

1. Marioni, R.E., McRae, A.F., Bressler, J. et al. Meta-analysis of epigenome-wide association studies of cognitive abilities. Mol Psychiatry 23, 2133–2144 (2018). 10.1038/s41380-017-0008-y

2. Smeeth D, Beck S, Karam EG, Pluess M. The role of epigenetics in psychological resilience. Lancet Psychiatry. 2021 Jul;8(7):620–629. doi: 10.1016/S2215-0366(20)30515-0. Epub 2021 Apr 27. PMID: 33915083; PMCID: PMC9561637

3. Deary, I.J., Cox, S.R. & Hill, W.D. Genetic variation, brain, and intelligence differences. Mol Psychiatry 27, 335–353 (2022). 10.1038/s41380-021-01027-y

4. Davies, et al. Study of 300,486 individuals identifies 148 independent genetic loci influencing general cognitive function. Nat Commun. 2018 May 29;9(1):2098. doi: 10.1038/s41467-018-04362-x.

5. Ross R, Goodpaster BH, Koch LG, Sarzynski MA, Kohrt WM, Johannsen NM, Skinner JS, Castro A, Irving BA, Noland RC, Sparks LM, Spielmann G, Day AG, Pitsch W, Hopkins WG, Bouchard C. Precision exercise medicine: understanding exercise response variability. Br J Sports Med. 2019 Sep;53(18):1141–1153. doi: 10.1136/bjsports-2018-100328. Epub 2019 Mar 12. PMID: 30862704; PMCID: PMC6818669.

6. Sanford, J.A., Nogiec, C.D., Lindholm, M.E., Adkins, J.N., Amar, D., Dasari, S., Drugan, J.K., Fernandez, F.M., Radom-Aizik, S., Schenk, S., Snyder, M.P., Tracy, R.P., Vanderboom, P., Trappe, S., Walsh, M.J., and the Molecular Transducers of Physical Activity Consortium (2020). Molecular transducers of physical activity consortium, (MoTrPAC): Mapping the dynamic responses to exercise. Cell 181, Cell 1464

7. Contrepois, K., Wu, S., Moneghetti, K.J., Hornburg, D., Ahadi, S., Tsai, M.-S.,, A.A., Wei, E., Lee-McMullen, B., Quijada, J.V., et al. (2020). Molecular Choreography of Acute Exercise. Cell 181, 1112–1130.e16. 10.1016/j.cell.2020.04.043.

8. Nair, V.D., Ge, Y., Li, S., Picas H., Jain N., Seenarine, N., Amper, M-A.S., Goodpaster, B.H., Walsh, M.J., Coen, P.M., Sealfon, S.C., 2020. Sedentary and trained older men have distinct circulating exosomal microRNA profiles at baseline and in response to acute exercise, Frontiers in Physiology, Vol. 11, Article 605

9. Marabita, F., James, T., Karhu, A., Virtanen, H., Kettunen, K., Stenlund, H., Boulund, F., Hellström, C., Neiman, M., Mills, R., et al. (2022). Multiomics and digital monitoring during lifestyle changes reveal independent dimensions of human biology and health. Cell Systems 13, 241–255.e7. 10.1016/j.cels.2021.11.001.

10. Watanabe, K., Wilmanski, T., Diener, C., Earls, J.C., Zimmer, A., Lincoln, B., Hadlock, J.J., Lovejoy, J.C., Gibbons, S.M., Magis, A.T., et al. (2023). Multiomic signatures of body mass index identify heterogeneous health phenotypes and responses to a lifestyle intervention. Nat Med 29, 996–1008. 10.1038/s41591-023-02248-0.

11. Franklin BA, Eijsvogels TMH, Pandey A, Quindry J, Toth PP. (2022) Physical activity, cardiorespiratory fitness, and cardiovascular health: A clinical practice statement of the American Society for Preventive Cardiology Part II: Physical activity, cardiorespiratory fitness, minimum and goal intensities for exercise training, prescriptive methods, and special patient populations. Am J Prev Cardiol. 2022 Oct 13;12:100425. doi: 10.1016/j.ajpc.2022.100425.

12. ACFT 2022. Army Combat Fitness Test https://www.army.mil/acft

13. PRT 2023: Navy Physical Readiness Test-https://www.mynavyhr.navy.mil/Portals/55/Support/Culture%20Resilience/Physical/Guide_5-Physical_Readiness_Test_PRT_JAN_2023.pdf?ver=OlmOLoZTfCA641JUkAnIaw%3D%3D

14. Bragazzi, N. L., Khoramipour, K., Chaouachi, A. & Chamari, K., (2020). Toward sportomics: Shifting from sport genomics to sport postgenomics and metabolomics specialties. Promises, challenges, and future perspectives. Int. J. Sports Physiol. Perform. 15, 1201–1202

15. McKetney, J., Jenkins, C.C., Minogue, C., Mach, P.M., Hussey, E.K., Glaros, T.G., Coon, J., and Dhummakupt, E.S. (2022). Proteomic and metabolomic profiling of acute and chronic stress events associated with military exercises. Mol. Omics 18, 279–295. 10.1039/D1MO00271F.

16. Bassini, A., Sartoretto, S., Jurisica, L., Magno-França, A., Anderson, L., Pearson, T., Razavi, M., Chandran, V., Martin III, L., Jurisica, I., Cameron, L.C. (2022), Sportomics method to assess acute phase proteins in Olympic level athletes using dried blood spots and multiplex assays. Nature, 12:19824.

17. Lombardo, B., Izzo, V., Terracciano, D., Ranieri, A., Mazzaccara, C., Fimiani, F., Cesaro, A., Gentile, L., Leggiero, E., Pero, R., et al. (2019). Laboratory medicine: health evaluation in elite athletes. Clinical Chemistry and Laboratory Medicine (CCLM) 57, 1450–1473. 10.1515/cclm-2018-1107.

18. Schwartz, P.L., Douglas, J.S., and Carroll, H.W. (1971). The Relationship Between Postexereise Concentration of Serum Pyruvate and Physical Fitness. Experimental Biology and Medicine 138, 130–136. 10.3181/00379727-138-35845.

19. Malsagova KA, Kopylov AT, Sinitsyna AA, Stepanov AA, Izotov AA, Butkova TV, Chingin K, Klyuchnikov MS, Kaysheva AL. Sports Nutrition: Diets, Selection Factors, Recommendations. Nutrients. 2021 Oct 25;13(11):3771. doi: 10.3390/nu13113771.

20. Pitsiladis, Y.P., Tanaka, M., Eynon, N., Bouchard, C., North, K.N., Williams, A.G., Collins, M., Moran, C.N., Britton, S.L., Fuku, N., et al. (2016). Athlome Project Consortium: a concerted effort to discover genomic and other “omic” markers of athletic performance. Physiol Genomics 48, 183–190. 10.1152/physiolgenomics.00105.2015.

21. Robbins, J.M., Peterson, B., Schranner, D., Tahir, U.A., Rienmüller, T., Deng, S., Keyes, M.J., Katz, D.H., Beltran, P.M.J., Barber, J.L., et al. (2021). Human plasma proteomic profiles indicative of cardiorespiratory fitness. Nat Metab 3, 786–797. 10.1038/s42255-021-00400-z.

22. Morville, T., Sahl, R.E., Moritz, T., Helge, J.W., and Clemmensen, C. (2020). Plasma Metabolome Profiling of Resistance Exercise and Endurance Exercise in Humans. Cell Reports 33, 108554. 10.1016/j.celrep.2020.108554.

23. Metwally, A.A., Zhang, T., Wu, S., Kellogg, R., Zhou, W., Contrepois, K., Tang, H., and Snyder, M. (2022). Robust identification of temporal biomarkers in longitudinal omics studies. Bioinformatics 38, 3802–3811. 10.1093/bioinformatics/btac403.

24. Ziyatdinov, A., Kim, J., Prokopenko, D., Privé, F., Laporte, F., Loh, P.-R., Kraft, P., and Aschard, H. (2021). Estimating the effective sample size in association studies of quantitative traits. G3 (Bethesda) 11, jkab057. 10.1093/g3journal/jkab057.

25. Peluso, A., Glen, R., and Ebbels, T.M.D. (2021). Multiple-testing correction in metabolome-wide association studies. BMC Bioinformatics 22, 67. 10.1186/s12859-021-03975-2.

26. DARPA MBA: Measuring Biological Aptitude https://www.darpa.mil/program/measuring-biological-aptitude.

27. Hardison, C.M., Mayberry, P.W., Krull, H., Setodji, C.M., Panis, C., Madison, R., Simpson, M., Avriette, M., Totten, M. (2022), Wong, J.; Independent Review of the Army Combat Fitness Test: Summary of Key Findings and Recommendations (RAND Corporation) 10.7249/RRA1825-1.

28. Army Publishing Directorate (2023): https://armypubs.army.mil/ProductMaps/PubForm/Details.aspx?PUB_ID=1026692

29. Szklarczyk, et al. (2019), STRING v11: protein-protein association networks with increased coverage, supporting functional discovery in genome-wide experimental datasets. Nucleic Acids Res., 47(D1): D607–D613

30. Kuhn M, Szklarczyk D, Pletscher-Frankild S, Blicher TH, von Mering C, Jensen LJ, Bork P. STITCH 4: integration of protein-chemical interactions with user data. Nucleic Acids Res. 2014 Jan;42(Database issue):D401–7. doi: 10.1093/nar/gkt1207.

31. Tuncbag, N., Gosline, S.J.C., Kedaigle, A., Soltis, A.R., Gitter, A., and Fraenkel, E. (2016). Network-Based Interpretation of Diverse High-Throughput Datasets through the Omics Integrator Software Package. PLoS Comput Biol 12, e1004879. 10.1371/journal.pcbi.1004879.

32. Federico A, Monti S (2020). “hypeR: an R package for geneset enrichment workflows.” Bioinformatics, 36(4), 1307–1308. R package version 1.14.0.

33. Mao W, Zaslavsky E, Hartmann BM, Sealfon SC, Chikina M. Pathway-level information extractor (PLIER) for gene expression data. Nat Methods. 2019 Jul;16(7):607–610. doi: 10.1038/s41592-019-0456-1. Epub 2019 Jun 27. PMID: 31249421; PMCID: PMC7262669.

34. Vandereyken K, Sifrim A, Thienpont B, Voet T. Methods and applications for single-cell and spatial multi-omics. Nat Rev Genet. 2023 Aug;24(8):494–515. doi: 10.1038/s41576-023-00580-2. Epub 2023 Mar 2. PMID: 36864178; PMCID: PMC9979144.

35. Roberts BM, Rushing KA, Plaisance EP. Sex Differences in Body Composition and Fitness Scores in Military Reserve Officers’ Training Corps Cadets. Mil Med. 2023 Jan 4;188(1-2):e1–e5. doi: 10.1093/milmed/usaa496. PMID: 33449115.

36. Boffey D, DiPrima JA, Kendall KL, Hill EC, Stout JR, Fukuda DH. Influence of Body Composition, Load-Velocity Profiles, and Sex-Related Differences on Army Combat Fitness Test Performance. J Strength Cond Res. 2023 Dec 1;37(12):2467–2476.

37. Sandhurst: https://www.westpoint.edu/sandhurst

38. Subramanian, A., Tamayo, P., Mootha, V.K., Mukherjee, S., Ebert, B.L., Gillette, M.A., Paulovich, A., Pomeroy, S.L., Golub, T.R., Lander, E.S., et al. (2005). Gene set enrichment analysis: A knowledge-based approach for interpreting genome-wide expression profiles. Proc. Natl. Acad. Sci. U.S.A. 102, 15545–15550. 10.1073/pnas.0506580102.

39. Barbie, D.A., Tamayo, P., Boehm, J.S., Kim, S.Y., Moody, S.E., Dunn, I.F., Schinzel, A.C., Sandy, P., Meylan, E., Scholl, C., et al. (2009). Systematic RNA interference reveals that oncogenic KRAS-driven cancers require TBK1. Nature 462, 108–112. 10.1038/nature08460.

40. West EE, Kolev M, Kemper C. Complement and the Regulation of T Cell Responses. Annu Rev Immunol. 2018 Apr 26;36:309–338. doi: 10.1146/annurev-immunol-042617-053245. PMID: 29677470; PMCID: PMC7478175

41. Rothschild-Rodriguez D, Causer AJ, Brown FF, Collier-Bain HD, Moore S, Murray J, Turner JE, Campbell JP. The effects of exercise on complement system proteins in humans: a systematic scoping review. Exerc Immunol Rev. 2022;28:1–35. PMID: 35452398.

42. Mastellos, D. C., Deangelis, R. A. & Lambris, J. D. Complement-triggered pathways orchestrate regenerative responses throughout phylogenesis. Semin. Immunol. 25, 29–38 (2013).

43. Chmilewsky, F., Jeanneau, C., Laurent, P., Kirschfink, M. & About, I. Pulp progenitor cell recruitment is selectively guided by a C5a gradient. J. Dent. Res. 92, 532–539 (2013).

44. Peterson, S. L., Nguyen, H. X., Mendez, O. A. & Anderson, A. J. Complement protein C1q modulates neurite outgrowth in vitro and spinal cord axon regeneration in vivo. J. Neurosci.: Off. J. Soc. Neurosci. 35, 4332–4349 (2015).

45. Marshall, K. M., He, S., Zhong, Z., Atkinson, C. & Tomlinson, S. Dissecting the complement pathway in hepatic injury and regeneration with a novel protective strategy. J. Exp. Med. 211, 1793–1805 (2014).

46. Zhang, C., Wang, C., Li, Y. et al. Complement C3a signaling facilitates skeletal muscle regeneration by regulating monocyte function and trafficking. Nat Commun 8, 2078 (2017). 10.1038/s41467-017-01526-z

47. Jia F, Liu X and Liu Y (2025) Bile acid signaling in skeletal muscle homeostasis: from molecular mechanisms to clinical applications. Front. Endocrinol. 16:1551100. doi: 10.3389/fendo.2025.1551100

48. Sasaki T, Kuboyama A, Mita M, Murata S, Shimizu M, Inoue J, Mori K, Sato R. The exercise-inducible bile acid receptor Tgr5 improves skeletal muscle function in mice. J Biol Chem. 2018 Jun 29;293(26):10322–10332. doi: 10.1074/jbc.RA118.002733. Epub 2018 May 17. PMID: 29773650; PMCID: PMC6028981.

49. Chen H, Ma J, Ma X. Administration of tauroursodeoxycholic acid attenuates dexamethasone-induced skeletal muscle atrophy. Biochem Biophys Res Commun. 2021 Sep 17;570:96–102. doi: 10.1016/j.bbrc.2021.06.102. Epub 2021 Jul 15. PMID: 34274852.

50. Viribay A, Burgos J, Fernández-Landa J, Seco-Calvo J, Mielgo-Ayuso J. Effects of Arginine Supplementation on Athletic Performance Based on Energy Metabolism: A Systematic Review and Meta-Analysis. Nutrients. 2020 May 2;12(5):1300. doi: 10.3390/nu12051300. PMID: 32370176; PMCID: PMC7282262.

51. Gonzalez AM, Townsend JR, Pinzone AG, Hoffman JR. Supplementation with Nitric Oxide Precursors for Strength Performance: A Review of the Current Literature. Nutrients. 2023 Jan 28;15(3):660. doi: 10.3390/nu15030660. PMID: 36771366; PMCID: PMC9921013.

52. Chen, S., Minegishi, Y., Hasumura, T. et al. Involvement of ammonia metabolism in the improvement of endurance performance by tea catechins in mice. Sci Rep 10, 6065 (2020

53. Lee EC, Fragala MS, Kavouras SA, Queen RM, Pryor JL, Casa DJ. Biomarkers in Sports and Exercise: Tracking Health, Performance, and Recovery in Athletes. J Strength Cond Res. 2017 Oct;31(10):2920–2937. doi: 10.1519/JSC.0000000000002122. PMID: 28737585; PMCID: PMC5640004.

54. Neill MA, Aschner J, Barr F, Summar ML. Quantitative RT-PCR comparison of the urea and nitric oxide cycle gene transcripts in adult human tissues. Mol Genet Metab. 2009 Jun;97(2):121–7. doi: 10.1016/j.ymgme.2009.02.009. Epub 2009 Mar 3. PMID: 19345634; PMCID: PMC2680466.

55. Cervenka I, Agudelo LZ, Ruas JL. (2017). Kynurenines: tryptophan’s metabolites in exercise, inflammation, and mental health. Science 357: eaaf9794, doi:10.1126/science.aaf9794.

56. Agudelo, L.Z., Femenía, T., Orhan, F., Porsmyr-Palmertz, M., Goiny, M., Martinez-Redondo, V., Correia, J.C., Izadi, M., Bhat, M., Schuppe-Koistinen, I., et al. (2014). Skeletal Muscle PGC-1α1 Modulates Kynurenine Metabolism and Mediates Resilience to Stress-Induced Depression. Cell 159, 33–45. 10.1016/j.cell.2014.07.051.

57. Morris, T.J., Butcher, L.M., Feber, A., Teschendorff, A.E., Chakravarthy, A.R., Wojdacz, T.K., and Beck, S. (2014). ChAMP: 450k Chip Analysis Methylation Pipeline. Bioinformatics 30, 428–430. 10.1093/bioinformatics/btt684.

58. Dobin, A., Davis, C.A., Schlesinger, F., Drenkow, J., Zaleski, C., Jha, S., Batut, P., Chaisson, M., and Gingeras, T.R. (2013). STAR: ultrafast universal RNA-seq aligner. Bioinformatics 29, 15–21. 10.1093/bioinformatics/bts635.

59. Love, M.I., Huber, W., and Anders, S. (2014). Moderated estimation of fold change and dispersion for RNA-seq data with DESeq2. Genome Biol 15, 550. 10.1186/s13059-014-0550-8.

60. Costa, P. T.; McCrae, R. R. (1976). Age differences in personality structure: A cluster analytic approach". Journal of Gerontology. 31 (5): 564–570.

61. Andrews. S., (2010) FastQC: A quality control tool for high throughput sequence data. URL: https://www.bioinformatics.babraham.ac.uk/projects/fastqc/.

62. Barcelo, H., Faul, J., Crimmins, E., Thyagarajan, B., (2018). A practical cryopreservation and staining protocol for immunophenotyping in population studies. Curr Protoc Cytom. April; 84(1): e35. doi:10.1002/cpcy.35.

63. Rohart F, Gautier B, Singh A, Lê Cao KA. mixOmics: An R package for ’omics feature selection and multiple data integration. PLoS Comput Biol. 2017 Nov 3;13(11)

64. Liberzon, A., Birger, C., Thorvaldsdóttir, H., Ghandi, M., Mesirov, J.P., and Tamayo, P. (2015). The Molecular Signatures Database Hallmark Gene Set Collection. Cell Systems 1, 417–425. 10.1016/j.cels.2015.12.004.

65. Wishart DS, Guo AC, Oler E, et al., HMDB 5.0: the Human Metabolome Database for 2022. Nucleic Acids Res. 2022. Jan 7;50(D1):D622–31.

