## Supplementary material for "Integration of Multiomic and Multi-phenotypic Data Identifies Biological Pathways Associated with Physical Fitness": multiple supplementary files: ACFT-PhenoMol-SUPPLMENTARY INFORMATION-Alizadeh2025JUL24.docx

Table ST1: Detailed description of molecular and cellular data used in the study (includes West Point male and female cohorts): BL = baseline prior to exercise, 2, 10 and 30 refer to the approximate number of minutes from cessation of exercise to blood draw. Bold cells indicate blood draws analyzed and included in analysis.

| **Study Segment** | **Subjects per Segment** | **Cellular Data Categories** | | **Molecular Data Categories** | | | | | |
| --- | --- | --- | --- | --- | --- | --- | --- | --- | --- |
|  |  | **Hematology**  (Fresh blood)  *38 markers/ draw* | **Immuno-phenotyping**  (PBMC)  *8 markers/ draw* | **iStat**  (Fresh blood)  *10 markers/ draw* | **3’ mRNA**  (Fresh blood)  *16318 transcripts/ draw* | **DNA Methyl-ation** (PBMC)  *16544 genes & 16415 promoters/ draw* | **Cytokines**  (Serum)  *14 markers/ draw* | **Untargeted Metabolome**  (Plasma)  *571 markers/ draw* | **Untargeted Proteome**  (Plasma)  *139 markers/ draw* |
| Segment 1 | 84 | **BL** | **BL, 30** | **BL, 2, 30** | **BL, 30** | **BL, 30** | **BL, 2, 30** | BL, 2, 30 | BL, 2, 30 |
| Segment 2 | 74 | **BL** | **BL, 30** | **BL, 2, 10, 30** | **BL** | **BL** | **BL, 2, 30** | **BL, 2, 10, 30** | **BL, 2, 10, 30** |
| Segment 3 | 60 | **BL** | **30** | **BL, 2, 10, 30** | **BL** | BL | BL, 2, 30 | BL, 2, 10, 30 | BL, 2, 10, 30 |
| Segment 4 | 57 | **BL** | **BL, 30** | **BL, 2, 10, 30** | **BL** | **BL, 30** | BL, 2, 30 | **BL, 2, 10, 30** | **BL, 2, 10, 30** |
| Segment 5 | 43 | **BL** | **BL, 30** | **BL, 2, 10, 30** | **BL, 30** | **BL, 30** | BL, 30 | **BL, 2, 10, 30** | **BL, 2, 10, 30** |
| Total time points measured/ analyzed | --- | **266** | **332** | **1187** | **420** | **338** | **115** | **696** | **693** |
| Total Molecular & Cellular measures used in the pipeline | ---- | **10,108** | **2,656** | **11,870** | **6,853,560** | **5,591,872 (genes)** | **1,610** | **397,416** | **96,327** |

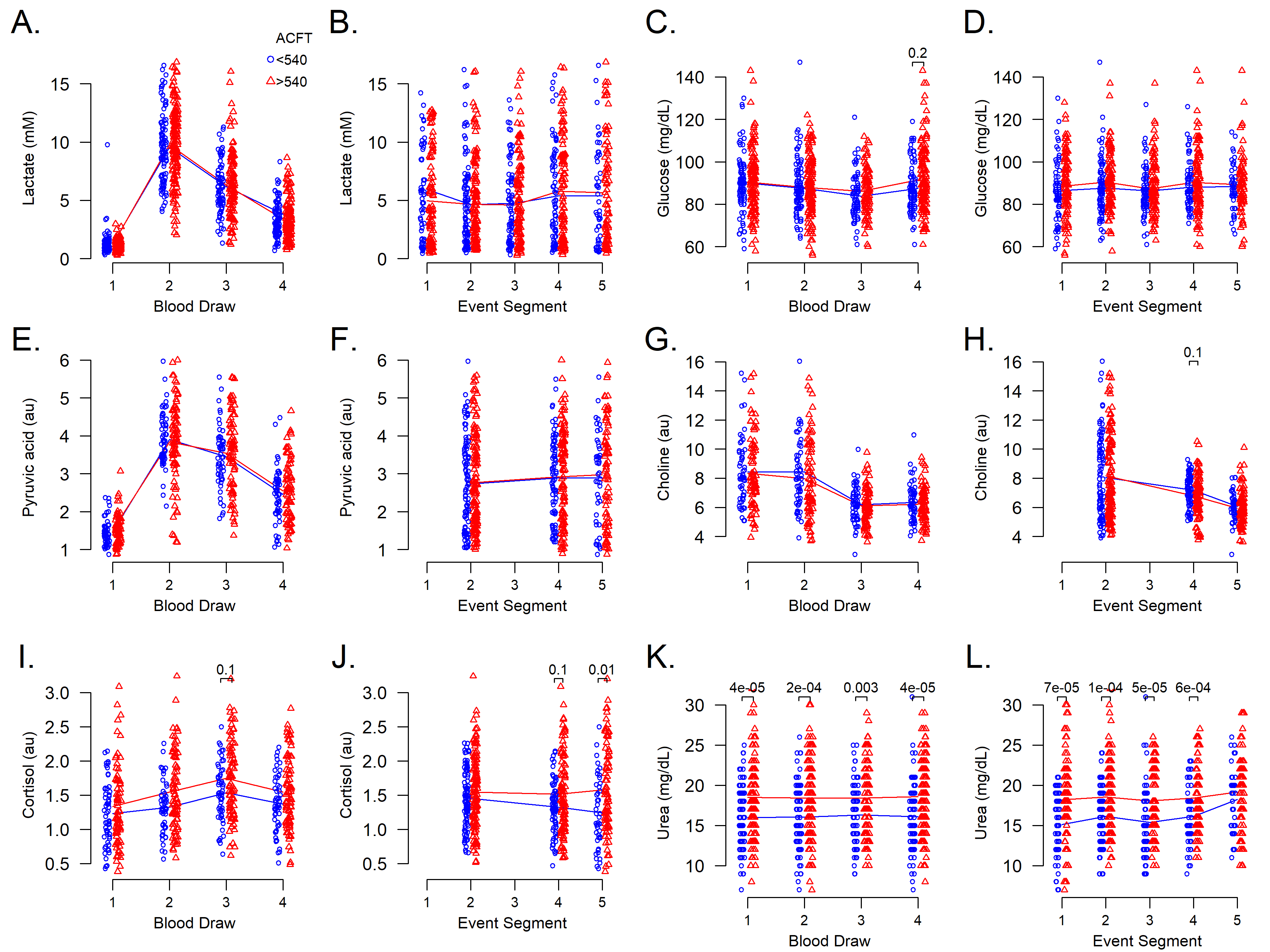

Figure S1: Short- and long-term longitudinal studies of blood biomarkers

Table ST28: Phenotype Feature Class

| **ACFT Event** | **Phenotype Name** | **Phenotype Group** | **Variable Contribution** | **Frequency Picked by sPLSR** | **Number Picked by sPLSR out of 30 analysis runs** | **Used in Pie Chart Summary** |
| --- | --- | --- | --- | --- | --- | --- |
| Maximum_Deadlift | LegToSkeletalMuscleMass | Body Composition | 812.4 | 100% | 30 | Yes |
| Maximum_Deadlift | BMI | Body Composition | 753.1 | 100% | 30 | Yes |
| Maximum_Deadlift | Body_Mass | Body Composition | 593.0 | 97% | 29 | Yes |
| Maximum_Deadlift | Skeletal_Muscle_Mass | Body Composition | 576.0 | 97% | 29 | Yes |
| Maximum_Deadlift | T4AvgPwr | O2 Utilization | 146.7 | 37% | 11 | No |
| Maximum_Deadlift | Time_To_Reach_VO2max_Minutes | O2 Utilization | 81.9 | 23% | 7 | No |
| Maximum_Deadlift | C5 | NEO-PI Conscientiousness | 14.7 | 3% | 1 | No |
| Maximum_Deadlift | Oura_Resting_Heart_Rate_Avg_bpm | Resting Heart Rate | 14.0 | 3% | 1 | No |
| Maximum_Deadlift | Leg_Lean_Mass | Body Composition | 8.3 | 3% | 1 | Yes |
| Standing_Power_Throw | Skeletal_Muscle_Mass | Body Composition | 705.1 | 100% | 30 | Yes |
| Standing_Power_Throw | Body_Mass | Body Composition | 567.8 | 100% | 30 | Yes |
| Standing_Power_Throw | Time_To_Reach_VO2max_Minutes | O2 Utilization | 561.1 | 100% | 30 | Yes |
| Standing_Power_Throw | LegToSkeletalMuscleMass | Body Composition | 489.1 | 90% | 27 | Yes |
| Standing_Power_Throw | BMI | Body Composition | 164.4 | 43% | 13 | Yes |
| Standing_Power_Throw | MCH_pg | Red Blood Cells | 146.2 | 37% | 11 | No |
| Standing_Power_Throw | MCV_fL | Red Blood Cells | 71.4 | 20% | 6 | No |
| Standing_Power_Throw | Oura_Avg_Resp_Rate_SD_brpm | Respiration Rate | 60.7 | 20% | 6 | No |
| Standing_Power_Throw | MCHC_Conc_g_dL | Red Blood Cells | 76.4 | 17% | 5 | No |
| Standing_Power_Throw | Leg_Lean_Mass | Body Composition | 50.8 | 13% | 4 | Yes |
| Standing_Power_Throw | WCST_PreStress_Conceptual_Level_Responses | WCST | 24.0 | 7% | 2 | No |
| Standing_Power_Throw | T4AvgPwr | O2 Utilization | 16.2 | 7% | 2 | Yes |
| Standing_Power_Throw | Cardio_Activity_Calories | Activity | 11.8 | 3% | 1 | No |
| Standing_Power_Throw | E2 | NEO-PI Extraversion | 11.6 | 3% | 1 | No |
| Standing_Power_Throw | NRBC_absNum_K_uL | Red Blood Cells | 11.4 | 3% | 1 | No |
| Standing_Power_Throw | HFLC_percent | White Blood Cells | 11.4 | 3% | 1 | No |
| Standing_Power_Throw | HFLC_absNum_K_uL | White Blood Cells | 10.7 | 3% | 1 | No |
| Standing_Power_Throw | MicroR_percent | Red Blood Cells | 9.8 | 3% | 1 | No |
| Hand_Release_Pushups | LegToSkeletalMuscleMass | Body Composition | 1020.0 | 87% | 26 | Yes |
| Hand_Release_Pushups | Time_To_Reach_VO2max_Minutes | O2 Utilization | 661.2 | 77% | 23 | Yes |
| Hand_Release_Pushups | NEUT_absNum_K_uL | White Blood Cells | 266.1 | 40% | 12 | No |
| Hand_Release_Pushups | Body_Height | Body Composition | 280.2 | 37% | 11 | Yes |
| Hand_Release_Pushups | NLR_Neutrophil_Lymphocyte_Ratio | White Blood Cells | 128.3 | 20% | 6 | No |
| Hand_Release_Pushups | O6 | NEO-PI Openness | 83.0 | 13% | 4 | No |
| Hand_Release_Pushups | WCST_PostStress_Nonperseverative_Errors_Score | WCST | 102.0 | 10% | 3 | No |
| Hand_Release_Pushups | Oura_Resting_Heart_Rate_Avg_bpm | Resting Heart Rate | 93.8 | 10% | 3 | No |
| Hand_Release_Pushups | C | NEO-PI Conscientiousness | 74.3 | 10% | 3 | No |
| Hand_Release_Pushups | C5 | NEO-PI Conscientiousness | 69.0 | 10% | 3 | No |
| Hand_Release_Pushups | C4 | NEO-PI Conscientiousness | 43.7 | 7% | 2 | No |
| Hand_Release_Pushups | WBC_absNum_K_uL | White Blood Cells | 33.5 | 7% | 2 | No |
| Hand_Release_Pushups | Number_of_Cardio_Activites | Activity | 26.8 | 3% | 1 | No |
| Hand_Release_Pushups | Average_Calories | Activity | 22.4 | 3% | 1 | No |
| Hand_Release_Pushups | BMI | Body Composition | 20.8 | 3% | 1 | Yes |
| Hand_Release_Pushups | N5 | NEO-PI Neuroticism | 20.5 | 3% | 1 | No |
| Hand_Release_Pushups | LYMPH_percent | White Blood Cells | 20.4 | 3% | 1 | No |
| Hand_Release_Pushups | Oura_Sleep_Time_SD_hrs | Sleep Variability | 18.4 | 3% | 1 | No |
| Hand_Release_Pushups | O | NEO-PI Openness | 15.7 | 3% | 1 | No |
| Sprint_Drag_Carry | Time_To_Reach_VO2max_Minutes | O2 Utilization | 976.7 | 100% | 30 | Yes |
| Sprint_Drag_Carry | T3AvgPwr | O2 Utilization | 634.5 | 87% | 26 | Yes |
| Sprint_Drag_Carry | LegToSkeletalMuscleMass | Body Composition | 369.8 | 53% | 16 | Yes |
| Sprint_Drag_Carry | Oura_Sleep_Time_SD_hrs | Sleep Variability | 297.8 | 50% | 15 | Yes |
| Sprint_Drag_Carry | Skeletal_Muscle_Mass | Body Composition | 194.3 | 30% | 9 | Yes |
| Sprint_Drag_Carry | Age_At_Enrollment | AGE | 101.3 | 17% | 5 | No |
| Sprint_Drag_Carry | Oura_Resting_Heart_Rate_Avg_bpm | Resting Heart Rate | 92.1 | 17% | 5 | No |
| Sprint_Drag_Carry | Oura_Deep_Sleep_SD_hrs | Sleep Variability | 57.9 | 13% | 4 | Yes |
| Sprint_Drag_Carry | Oura_Avg_Resp_Rate_SD_brpm | Respiration Rate | 55.9 | 13% | 4 | No |
| Sprint_Drag_Carry | T4AvgPwr | O2 Utilization | 36.7 | 7% | 2 | Yes |
| Sprint_Drag_Carry | Body_Mass | Body Composition | 25.5 | 7% | 2 | Yes |
| Sprint_Drag_Carry | O6 | NEO-PI Openness | 37.6 | 3% | 1 | No |
| Sprint_Drag_Carry | MCH_pg | Red Blood Cells | 28.0 | 3% | 1 | No |
| Sprint_Drag_Carry | STANINE_Raven_Progressive_Matrix | RPM Test | 24.3 | 3% | 1 | No |
| Sprint_Drag_Carry | TNAscore_Raven_Progressive_Matrix | RPM Test | 20.9 | 3% | 1 | No |
| Sprint_Drag_Carry | Correct_Raven_Progressive_Matrix | RPM Test | 16.1 | 3% | 1 | No |
| Sprint_Drag_Carry | BMI | Body Composition | 16.0 | 3% | 1 | Yes |
| Sprint_Drag_Carry | MCV_fL | Red Blood Cells | 14.8 | 3% | 1 | No |
| Leg_Tuck_OR_Plank | VO2_Max_Bike | O2 Utilization | 863.7 | 93% | 28 | Yes |
| Leg_Tuck_OR_Plank | Cardio_Activity_Avg_Calories | Activity | 553.7 | 63% | 19 | Yes |
| Leg_Tuck_OR_Plank | Oura_Sleep_Time_SD_hrs | Sleep Variability | 430.5 | 57% | 17 | Yes |
| Leg_Tuck_OR_Plank | Average_Calories | Activity | 224.2 | 40% | 12 | Yes |
| Leg_Tuck_OR_Plank | Time_To_Reach_VO2max_Minutes | O2 Utilization | 194.5 | 33% | 10 | Yes |
| Leg_Tuck_OR_Plank | LegToSkeletalMuscleMass | Body Composition | 182.6 | 30% | 9 | No |
| Leg_Tuck_OR_Plank | O5 | NEO-PI Openness | 119.8 | 20% | 6 | No |
| Leg_Tuck_OR_Plank | T3AvgPwr | O2 Utilization | 48.3 | 13% | 4 | Yes |
| Leg_Tuck_OR_Plank | Oura_Avg_Resp_Rate_SD_brpm | Respiration Rate | 89.9 | 10% | 3 | No |
| Leg_Tuck_OR_Plank | Oura_Deep_Sleep_SD_hrs | Sleep Variability | 40.9 | 10% | 3 | Yes |
| Leg_Tuck_OR_Plank | T2AvgHR | O2 Utilization | 60.3 | 7% | 2 | Yes |
| Leg_Tuck_OR_Plank | Oura_Resting_Heart_Rate_Avg_bpm | Resting Heart Rate | 50.5 | 7% | 2 | No |
| Leg_Tuck_OR_Plank | Body_Fat_Percent | Body Composition | 44.4 | 7% | 2 | No |
| Leg_Tuck_OR_Plank | N5 | NEO-PI Neuroticism | 26.1 | 7% | 2 | No |
| Leg_Tuck_OR_Plank | E1 | NEO-PI Extraversion | 23.4 | 3% | 1 | No |
| Leg_Tuck_OR_Plank | Oura_Awake_Sleep_SD_hrs | Sleep Variability | 16.3 | 3% | 1 | Yes |
| Leg_Tuck_OR_Plank | Number_of_Cardio_Activites | Activity | 16.0 | 3% | 1 | Yes |
| Leg_Tuck_OR_Plank | Body_Height | Body Composition | 15.0 | 3% | 1 | No |
| 2_Mile_Run | VO2_Max_Bike | O2 Utilization | 786.3 | 80% | 24 | Yes |
| 2_Mile_Run | T3AvgPwr | O2 Utilization | 374.2 | 53% | 16 | Yes |
| 2_Mile_Run | Oura_REM_Sleep_SD_hrs | Sleep Variability | 338.2 | 47% | 14 | Yes |
| 2_Mile_Run | Oura_Sleep_Time_SD_hrs | Sleep Variability | 218.3 | 37% | 11 | Yes |
| 2_Mile_Run | Oura_Resting_Heart_Rate_Avg_bpm | Resting Heart Rate | 374.9 | 30% | 9 | No |
| 2_Mile_Run | T4AvgPwr | O2 Utilization | 191.9 | 30% | 9 | Yes |
| 2_Mile_Run | Body_Fat_Percent | Body Composition | 157.8 | 27% | 8 | No |
| 2_Mile_Run | O2 | NEO-PI Openness | 137.4 | 27% | 8 | No |
| 2_Mile_Run | O5 | NEO-PI Openness | 139.5 | 20% | 6 | No |
| 2_Mile_Run | P_LCR_percent | Platelets | 90.9 | 13% | 4 | No |
| 2_Mile_Run | Oura_Deep_Sleep_SD_hrs | Sleep Variability | 54.1 | 7% | 2 | Yes |
| 2_Mile_Run | MPV_fL | Platelets | 39.1 | 7% | 2 | No |
| 2_Mile_Run | PDW_fL | Platelets | 35.9 | 7% | 2 | No |
| 2_Mile_Run | Average_Calories | Activity | 20.4 | 3% | 1 | No |
| 2_Mile_Run | Oura_Avg_Resp_Rate_SD_brpm | Respiration Rate | 15.9 | 3% | 1 | No |
| 2_Mile_Run | WBC_absNum_K_uL | White Blood Cells | 13.0 | 3% | 1 | No |
| 2_Mile_Run | Cardio_Activity_Calories | Activity | 12.0 | 3% | 1 | No |
